# Modeling the Multiple Facets of Speciation-with-Gene-Flow Towards Inferring the Divergence History of Lake Whitefish Species Pairs (Coregonus Clupeaformis)

**DOI:** 10.1101/068932

**Authors:** Clément Rougeux, Louis Bernatchez, Pierre-Alexandre Gagnaire

## Abstract

Parallel divergence patterns across replicated species pairs occurring in similar environmental contrasts may arise through distinct evolutionary scenarios. Deciphering whether such parallelism actually reflects repeated parallel divergence driven by divergent selection or a single divergence event with subsequent gene flow needs to be ascertained. Reconstructing historical gene flow is therefore of fundamental interest to understand how demography and selection jointly shaped genomic divergence during speciation. Here, we use an extended modeling framework to explore the multiple facets of speciation-with-gene-flow with demo-genetic divergence models that capture both temporal and genomic variation in effective population size and migration rate. We investigate the divergence history of five sympatric Lake Whitefish limnetic (dwarf) and benthic (normal) species pairs characterized by variable degrees of ecological divergence and reproductive isolation. Genome-wide SNPs were used to document the extent of genetic differentiation in each species pair, and 26 divergence models were fitted and compared to the unfolded joint allele frequency spectrum of each pair. We found evidence that a recent (circa 3000-4000 generations) asymmetrical secondary contact between expanding post-glacial populations has accompanied Whitefish diversification. Our results suggest that heterogeneous genomic differentiation patterns have emerged through the combined effects of linked selection generating variable rates of lineage sorting across the genome during geographical isolation, and heterogeneous introgression eroding divergence at different rates across the genome upon secondary contact. This study thus provides a new retrospective insight into the historical demographic and selective processes that shaped a continuum of divergence associated with ecological speciation.

## Introduction

Historical changes in the geographical distribution of species have been an important driver of diversification across many taxa (Coyne and Orr 2004). In particular, the pronounced climatic variations that occurred during the late Pleistocene caused major shifts in the distribution ranges of many species. These shifts are responsible for the divergence of ancestral lineages that survived in different glacial refugia, and then possibly came into secondary contact during interglacial periods (Bernatchez and Wilson 1998; Avise 2000; Hewitt 2001). The signature of post-glacial recolonization is still apparent in well-known terrestrial and aquatic suture zones, where multiple contacts between expanding post-glacial lineages tend to overlap and form hybrid zones hotspots for many species (Hewitt 1996; Hewitt 2000; Hewitt 2004; Swenson and Howard 2005; Bierne et al. 2011; April et al. 2013).

In some cases, secondary contacts have resulted in the sympatric enclosure of previously allopatric, partially reproductively isolated lineages, for instance within post-glacial lakes. This sympatric coexistence should have facilitated gene flow compared to parapatric populations, eventually leading to complete genetic homogenization of the original glacial lineages. This is not the case, however, for several north temperate freshwater fishes in which sympatric glacial lineages have further diverged into phenotypically differentiated and reproductively isolated species pairs following secondary contact (Bernatchez and Dodson 1990; McPhail 1992; Taylor and Bentzen 1993; Schluter 1996; Wood and Foote 1996; Taylor 1999). These cases of ecological speciation have been hypothesized to reflect adaptive responses to minimize competitive interactions and outbreeding depression through ecological niche segregation and hybridization avoidance among previously allopatric lineages (Bernatchez et al. 2010).

The genomic processes responsible for the phenotypic diversification of these incipient sympatric species remain contentious, especially with regards to the relative contributions of genetic differences that evolved in allopatry compared to more recent genetic changes occurring in sympatry (Welch and Jiggins 2014). To gain a more thorough understanding of how divergence unfolds at the molecular level, it is crucial to simultaneously take into account the historical demographic events that accompanied divergence and the subsequent genetic exchanges that occurred in sympatry. Genome-wide polymorphism data now provide the opportunity to infer complex demographic histories (Gutenkunst et al. 2009; Excoffier et al. 2013; Butlin et al. 2014) and investigate the evolutionary processes leading to the formation of nascent sympatric species.

Many aspects of populations' evolutionary history are influenced by demography, such as the rate of lineage sorting and gene exchange (Sousa and Hey 2013). Several approaches have been developed to infer the history of population divergence from genetic data obtained from contemporary populations. These methods usually rely on demographic models capturing the effects of population size, splitting time and migration between two populations exchanging genes (Hey and Nielsen 2004; Becquet and Przeworski 2007; Hey and Nielsen 2007). An important facet of the speciation process which is usually not taken into account by demographic models is that a significant proportion of the genome may be affected by selection (Barton and Bengtsson 1986; Feder et al. 2012; Nosil 2012; Harrison and Larson 2016; Wolf and Ellegren 20161). Two different selective processes that generate heterogeneous genome divergence can be distinguished. One occurs during contact episodes and corresponds to selection acting on genomic regions involved in reproductive isolation and local adaptation (Harrison 1990; Wu 2001; Payseur 2010). The second occurs through the action of background selection and selective sweeps (Hill and Robertson 1966; Smith and Haigh 1974; Charlesworth et al. 1997), which remove linked neutral diversity within populations during periods of reduced gene flow (Nachman and Payseur 2011; Cruickshank and Hahn 2014). While the barrier effect of speciation genes is equivalent to a local reduction in effective migration rate (m_e_) (Barton and Bengtsson 1986; Feder and Nosil 2010), linked selection rather corresponds to a reduction in effective population size (N_e_) that locally accelerates lineage sorting in the genome (Charlesworth 2006). Based on this, it is possible to capture the barrier effect of speciation genes by allowing for varying rates of introgression among loci (Roux et al. 2013; Sousa and Hey 2013; Tine et al. 2014), and to capture the effect of linked selection by allowing loci to experience varying rates of genetic drift (Sousa and Hey 2013; Roux et al. 2016). This provides a framework in which the effects of heterogeneous selection and gene flow can be considered separately or simultaneously to identify the simplest divergence scenario that best explain the data, thus avoiding overparametrization issues.

North American Lake Whitefish *(Coregonus clupeaformis)* represents a valuable model to study the role of past allopatric isolation on recent, sympatric ecological divergence. The Saint-John River drainage (southeastern Québec, northeastern Maine), where benthic (normal) and limnetic (dwarf) Whitefish sympatric species pairs occur in different lakes, corresponds to a suture zone where two glacial lineages (Atlantic/Mississippian and Acadian) have been hypothesized to have come into secondary contact during the last glacial retreat on the basis of mitochondrial DNA phylogeography (Bernatchez and Dodson 1990). Given, the historical hydrology of the region whereby there was a limited temporal window during which different lakes could be colonized by fish before becoming isolated (Curry 2007), the most likely scenario is that of an independent phenotypic divergence that occurred in each lake (Bernatchez 2004). In some lakes, phenotypic divergence between sympatric dwarf (limnetic) and normal (benthic) populations is still partly associated with the mitochondrial DNA lineages characterizing the different glacial origins of the sympatric populations (Pigeon et al. 1997). Dwarf whitefish are most often associated with the Acadian mitochondrial lineage and are only found in sympatry with the normal species. Moreover, fish from the Acadian lineage have a normal phenotype outside the contact zone, which supports the hypothesis that the dwarf species has been derived postglacially from an Acadian genetic background within the contact zone and independently in each lake (Bernatchez and Dodson 1990; Bernatchez and Dodson 1991; Bernatchez et al. 2010). The evolution of further phenotypic divergence in sympatry suggests that character displacement (Bernatchez 2004) may have been facilitated by the contact between genetically differentiated lineages. Moreover, the different dwarf-normal species pairs found in the contact zone are arrayed along a continuum of phenotypic differentiation, which closely mirrors the potential for niche segregation and exclusive interactions within lakes (Lu and Bernatchez 1999; Rogers et al. 2002; Landry et al. 2007; Rogers and Bernatchez 2007; Landry and Bernatchez 2010). This continuum is also evident at the genomic level, with increased baseline genetic differentiation and larger genomic islands of differentiation being found from the least to the most phenotypically and ecologically differentiated species pair (Renaut et al. 2012; Gagnaire et al. 2013b). Finally, quantitative trait loci (QTL) underlying adaptive phenotypic divergence map preferentially to genomic islands of differentiation (Renaut et al. 2012; Gagnaire et al. 2013a, 2013b) suggesting that selection acting on these traits contribute to the barrier to gene flow. Despite such detailed knowledge on this system, previous studies did not allow clarifying how the genomic landscape of dwarf-normal divergence in each lake has been influenced by the relative effects of directional selection on these QTLs and post-glacial differential introgression. Consequently, it is fundamental to elucidate the demographic history of the dwarf-normal whitefish species pairs to disentangle the evolutionary mechanisms involved in their diversification.

The main goal of this study was to use a genome-wide single nucleotide polymorphism (SNP) dataset to infer the demographic history associated with the recent phenotypic diversification of five sympatric dwarf and normal Lake Whitefish species pairs. Using RAD-seq SNP data to document the Joint Allele Frequency Spectrum (JAFS) in each species pair, we specifically test for the role of temporal and genomic variations in the rate of gene flow for each species pair separately, controlling for both effective population size and migration. We then performed historical gene flow analyses among lakes to determine how the different scenarios independently inferred within each lake collectively depict a parsimonious evolutionary scenario of diversification. Finally, we document how the complex interplay between historical contingency, demography and selection jointly shaped the continuum of divergence among sympatric whitefish species pairs.

## Results

### Comparisons among divergence models

For each of the five sympatric whitefish species pairs studied (Fig. 1), 26 alternative divergence models of increasing complexity were fitted to polymorphism data and compared to each other. This comparative framework enabled us to account for four different aspects of gene flow, that were considered separately or in combination. This includes temporal variation in effective population size *(N_e_)* and migration rate (m), but also genomic heterogeneity in these parameters to capture selective effects (Fig. 2). The unfolded JAFS of each species pair was constructed using orientated SNPs for which we could distinguish the ancestral and the derived variant. The five JAFS obtained highlighted the continuum of divergence existing among lakes (Fig. 3A, Supplementary Fig. S1). Namely, the density of shared polymorphisms located along the diagonal decreased from the least divergent (Témiscouata, East and Webster) to the most divergent (Indian and Cliff) species pairs, while the variance of SNP density around the diagonal increased accordingly. In addition, non-shared polymorphisms *(i.e.,* private SNPs) located on the outer frame of the spectrum were mostly found in Indian and Cliff lakes, which were also the only lakes to display differentially fixed SNPs between dwarf and normal whitefish *(i.e.,* F_st_=1).

**Figure 1:**
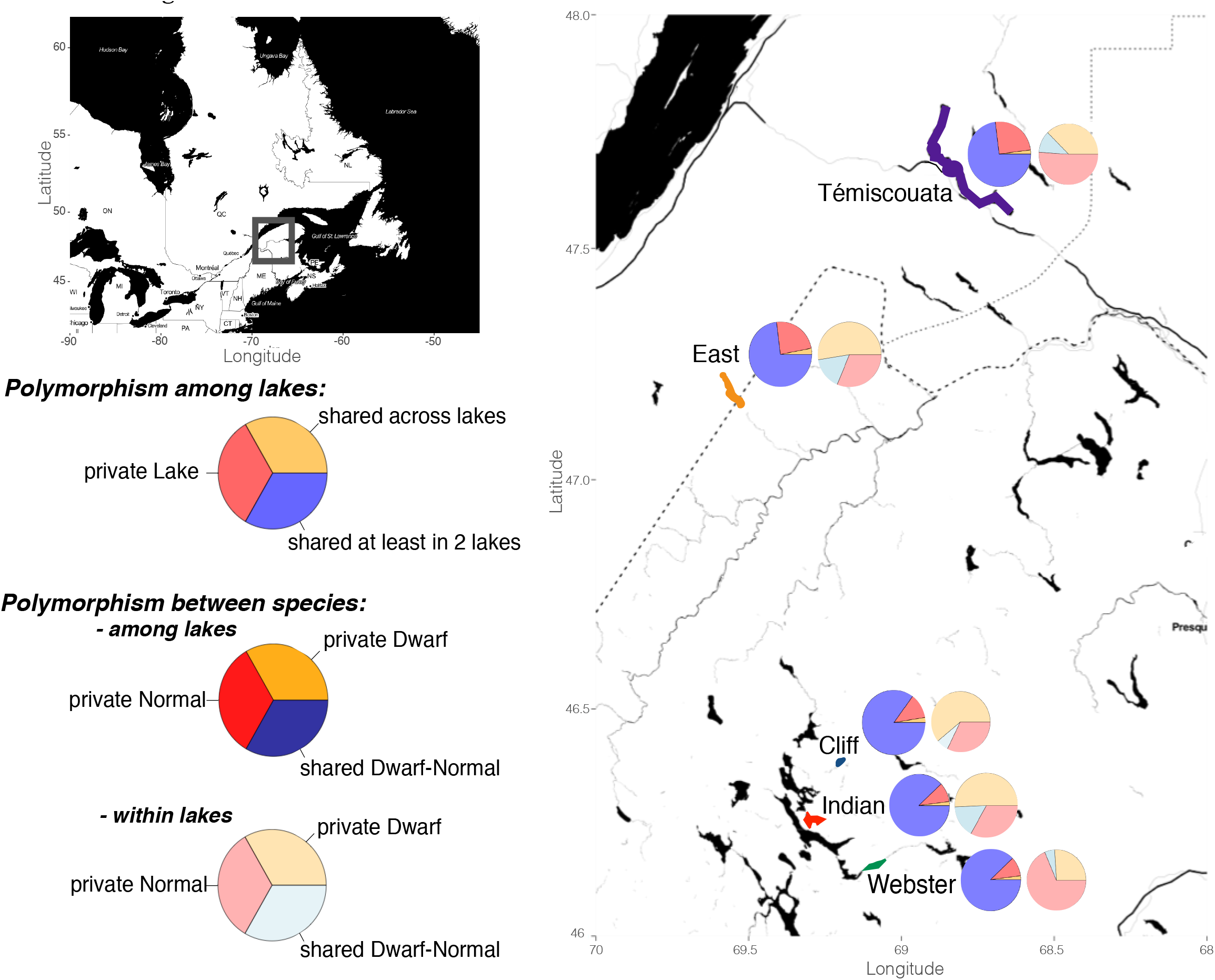
Geographic locations of lakes where sympatric lake whitefish species pairs were sampled, and overview of the extent of shared polymorphism. Each pie chart corresponds to the shared and private polymorphism among lakes and between species ecotypes. The first pie chart documents the sharing of polymorphism among lakes by distinguishing three categories of SNPs: 'private to a lake', alleles 'shared in at least two lakes' and 'shared across all lakes'. The other pie chart aims to describe the sharing of polymorphism between species at two levels; among lakes and within lakes, where SNPs were categorized as private to a species or shared between species

**Figure 2:**
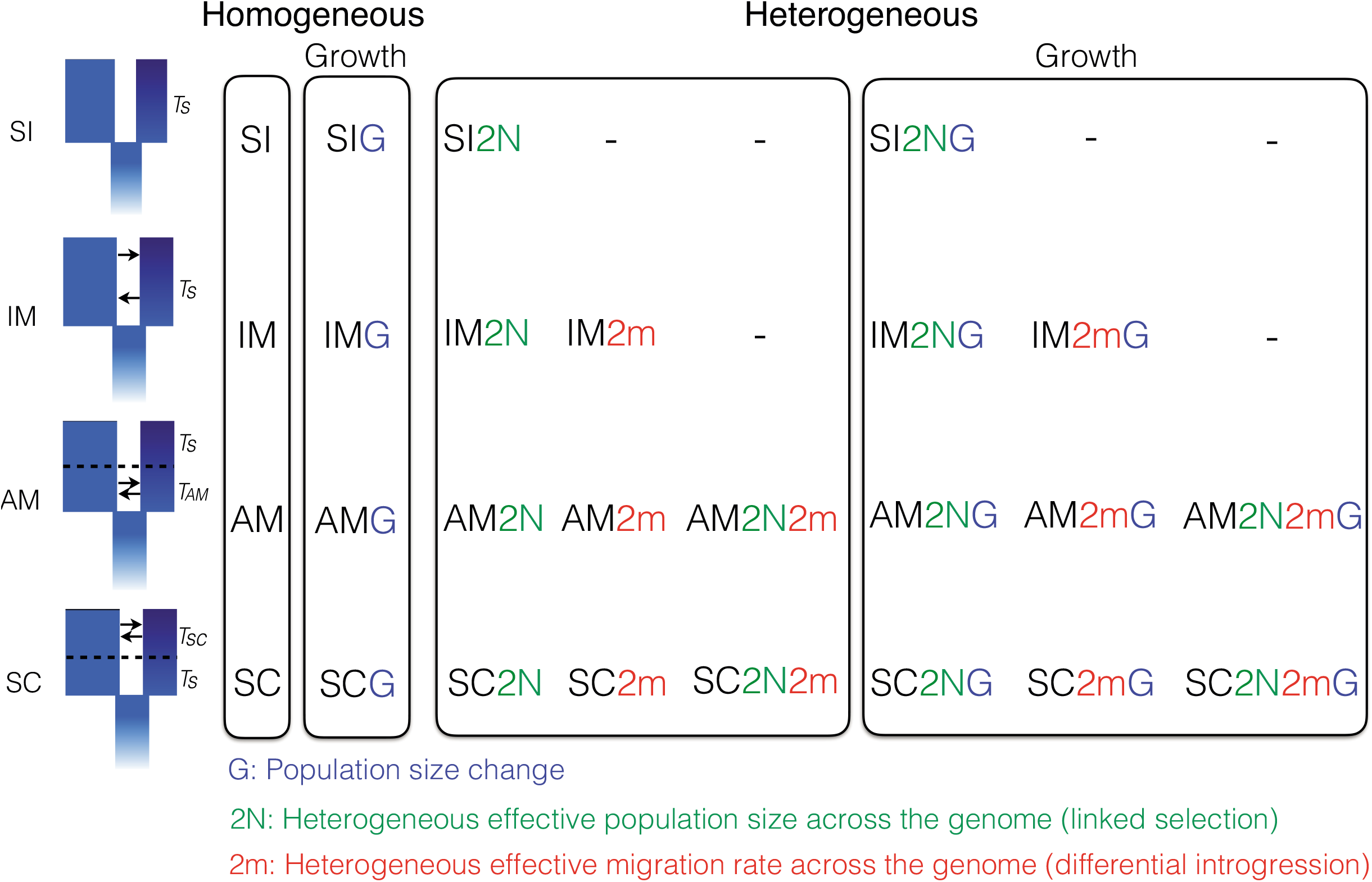
The 26 models implemented in the study. All the models implemented in this study were based on the four classical models of divergence: 'Strict Isolation' (-SI), 'Isolation with Migration' (-IM), 'Ancient Migration' (-AM) and 'Secondary Contact' (-SC). Briefly, *Ts* corresponds to the duration of complete isolation between diverging populations and *TAM* and *TSC* correspond to the duration of gene-flow in AM and SC models, respectively. The first extension of these four models accounts for temporal variation in effective populations size (-G models), allowing independent expansion or reduction of the diverging populations. The last categories of models correspond the 'Heterogeneous gene flow' models, which integrate parameters allowing genomic variations in migration rate (−2m), effective population size (−2N) or both (−2m2N) to account for genetic barriers and selection at linked sites.

**Figure 3:**
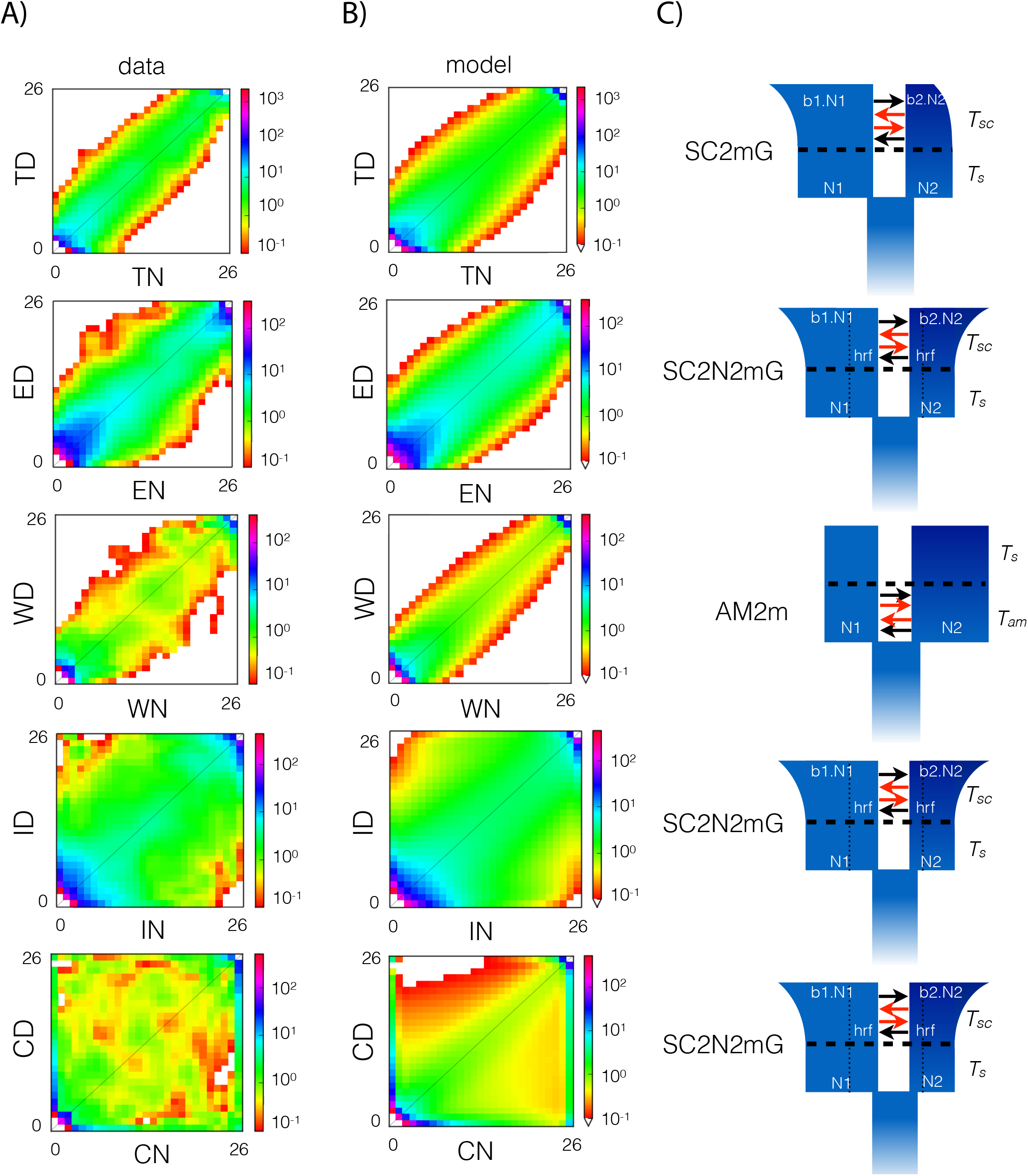
Historical demography of the Lake Whitefish species pairs. **(A)** Observed joint allele frequency spectrum (JAFS) for normal (-N; x-axis) and dwarf (-D; y-axis) populations for each lake (T=Témiscouata, E=East, W=Webster, I=Indian and C=Cliff), obtained by projection of empirical data. For each JAFS, the color scale corresponds to the probability of occurrence of the derived allele from 13 individuals from each population. **(B)** Predicted JAFS of the fittest model per lake. **(C)** Representation of the fittest model corresponding for each lake.

The comparison of model scores among lakes showed the importance of considering temporal changes in effective population size. Models including population growth (-G models) generally provided better fits to the data for Témiscouata, Indian and Cliff lakes (Mann-Whitney U test, *p* = 0.002; *p* = 0.011 and *p* = 0.0001, Fig. 4A). Similarly, accounting for heterogeneous migration rates across the genome (-2m models) improved the average model scores for each lake, although not significantly in East and Webster lakes (U test, *p* = 0.016; *p* = 0.063 and *p* = 0.029 for Témiscouata, Indian and Cliff lakes, respectively) (Fig. 4B). Moreover, models integrating heterogeneous effective population size at the genomic level (-2N models) provided significant improvements for the three most divergent species pairs Webster, Indian and Cliff lakes (U test, *p* = 0.013; *p* = 0.010 and *p* = 0.005, respectively) (Fig. 4C).

**Figure 4:**
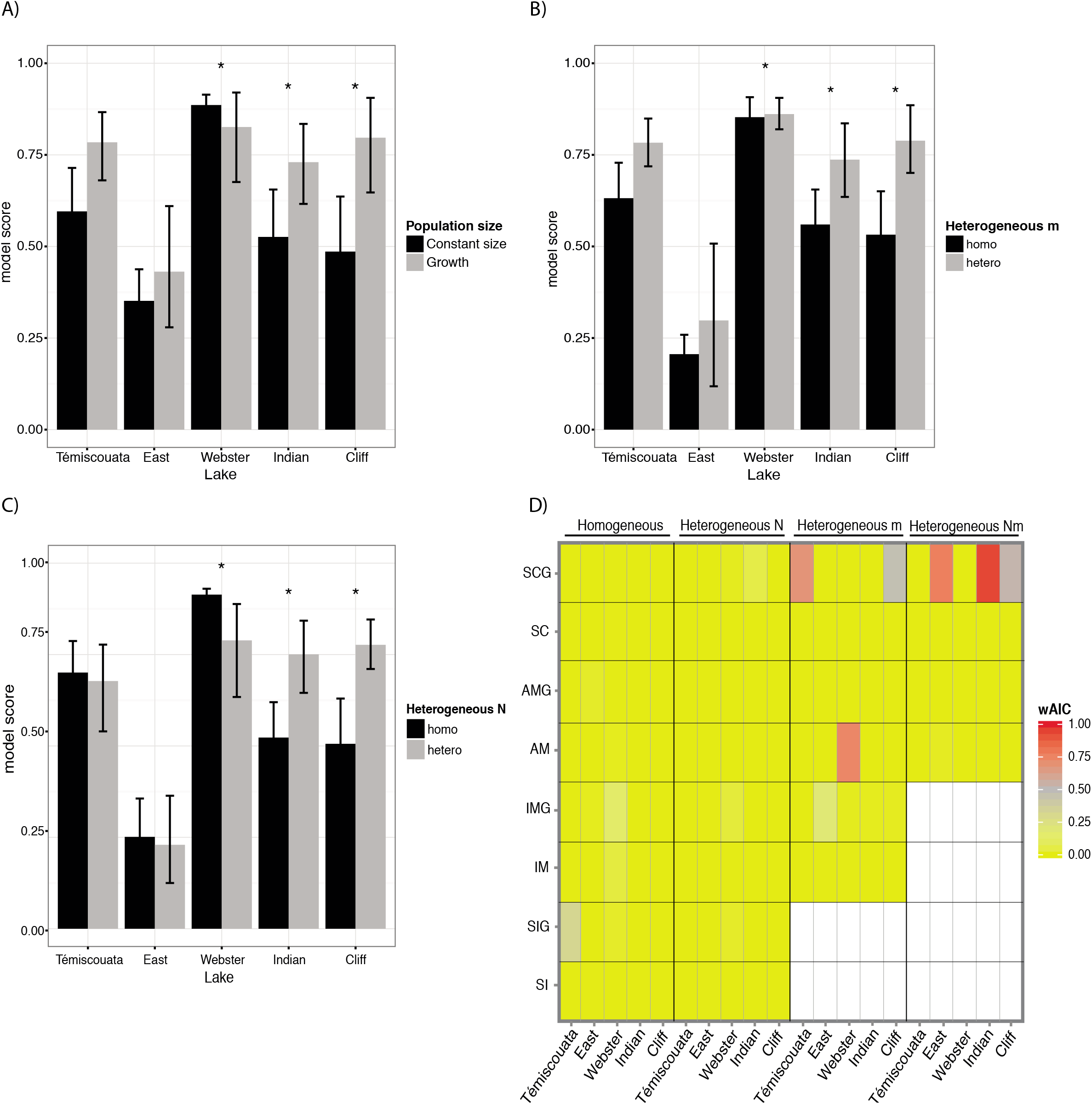
Parameters effects on inferred models and comparisons among models. Barplots showing the effect of taking into account particular demographic or selective aspects in the models, assessed using model scores, with **(A)** the effect of including temporal variation in population effective size (-G), **(B)** heterogeneous migration rates among loci (−2m) and **(C)** heterogeneous effective population sizes among loci (−2N). The vertical bars indicate the variance of the model scores within a model category (e.g. with or without -G parameter) and asterisks represent significant differences in average model scores between the compared categories of models. **(D)** Heat-map of the weighted AIC (w_AIC_) showing the relative weights of the 26 models for each lake. The color scale corresponds to the *w_AIC_* values ranging from 0 to 1. Warmer colors indicate the best models for each lake.

We performed model selection based on the AIC to penalize model likelihood by the number of parameters to avoid over-fitting. Applying a criterion of *ΔAIC_i_ ≤* 10 for model selection, we retained two best models for Témiscouata (SC2mG and SIG), Indian (SC2N2mG and SC2NG) and Cliff (SC2N2mG and SC2mG) lakes, four best models for East Lake (SC2N2mG, IM2mG, AM2N2m, AMG), and eight models for Webster Lake (AM2m, IMG, IM, IM2NG, SIG, SI2NG, IM2mG and SC2NG) (Table 1, Supplementary Table S1). The high number of retained models for Webster illustrates the difficulty to distinguish among different scenarios in this lake, which also displayed a particularly reduced level of polymorphism compared to the four other lakes (Fig. 3A). Akaike weights (w_AIC_) were higher than 0.5 for the highest ranked model of each lake (Fig. 4D, Table 1, Supplementary Table S1). For Témiscouata, the best model was a secondary contact with heterogeneous migration contemporary to population size change (SC2mG) which had a weight of 0.68. The best model for Webster Lake was a scenario of ancient migration with heterogeneous migration (AM2m, *w_AIC_ =* 0.74). Finally, East Lake and the two most differentiated species pairs (Indian and Cliff) received the highest support for a secondary contact model with heterogeneous migration rate and effective population size contemporary to population size change (SC2N2mG; *w_AIC_ =* 0.77 in East, *w_AIC_ =* 0.94 in Indian and *w_AIC_ =* 0.53 in Cliff).

**Table 1.** **Converted model parameter values for the best fit models in each lake.** For all lakes, statistics and demographic parameters details of the fittest models under the fixed threshold of Δ*AlC_t_ <* 10. The table contains in order the maximum likelihood (MLE) obtained from model with the smallest Akaike information criterion (AIC), the Δ*AI*C value for the corresponding model and the weighted AIC (w_AIC_). Then, the inferred demographic parameters values scaled by Theta *(Θ)*: the ancestral effective population size before population split (Nref); the effective population size after split for dwarf *(N_1_)* and normal (N_2_) populations; the growth coefficient for dwarf (b_1_) and normal (b_2_) populations. The *b* parameter defined as a ratio of contemporary effective population size of the ancestral populations at time of split (models: SI, IM and AM) and at time of secondary contact (models SC). Population exponential growth is associated with b_i_>1 and reduction in population effective size with bi<1; the Hill-Robertson factor (hrf) which corresponds to the degree to which the effective population size of the diverging populations (not considering the ancestral population) is locally reduced due to the effect of background selection or selective sweep effects; migrations rates from normal population to dwarf population *(m_e12_)* and reciprocally *(m_e21_);* the effective migration rates for genomic regions under selection (m_e12_) and *(m_e_'_21_);* temporal estimates (in years) regarding the split of the ancestral population (Tsplit) and the time when the migration stopped *(i.e., T_AM_* for the AM based models) and started after secondary contact *(i.e., T_SC_).* Finally, the table also contains proportion parameters such as the proportion *(Q)* of the genome with a reduced effective population size due to selection at linked sites; the proportion *(P)* of the genome evolving neutrally and the proportion *(O)* of accurate orientation of our SNPs. The estimation of each parameter in each case was translated in number of migrant/generation for migrations rates and temporal parameter in years, for a better understanding. Number in brackets denote 95% confidence interval obtained by 1000 bootstrap over loci.

| LAKE | MODEL | k | MLE | AIC | ΔAIC | wAIC | $\theta$ | Nref | $N_1$ | $N_2$ | $b_1$ | $b_i$ |
| --- | --- | --- | --- | --- | --- | --- | --- | --- | --- | --- | --- | --- |
| Témiscouata | SC2mG | 12 | -1656.47 | 3336.93 | - | 0.68 | 532.15 | 7611.39 | 2952603.4 | 1120233.2 | 20.3 | 0.3 |
|  |  |  |  |  |  |  |  |  | [870315.0; 5034891.7] | [266.4; 2253298.1] | [0.0; 68.6] | [0.0; 32.4] |
| Témiscouata | SIG | 6 | -1663.23 | 3338.47 | 1.54 | 0.32 | 570.56 | 4350.13 | 507815.2 | 749808.8 | 95.4 | 0.2 |
|  |  |  |  |  |  |  |  |  | [330990.9; 684639.4] | [717244.8; 782372.8] | [81.0; 109.8] | [0.0; 1.5] |
| East | SC2N2mG | 14 | -909.87 | 1847.74 | - | 0.77 | 212.86 | 1622.96 | 2325.3 | 2453.9 | 37.0 | 2.2 |
|  |  |  |  |  |  |  |  |  | [56.8; 54009.3] | [56.8; 38360.6] | [0.0; 104.7] | [0.0; 41.1] |
| East | IM2mG | 11 | -914.25 | 1850.50 | 2.77 | 0.19 | 982.85 | 7493.63 | 27310.7 | 2249.4 | 1.8 | 84.9 |
|  |  |  |  |  |  |  |  |  | [262.3; 57150.4] | [262.3; 30703.2] | [0.0; 3.7] | [82.1; 87.7] |
| East | AM2N2m | 12 | -916.30 | 1856.60 | 7.71 | 0.02 | 406.95 | 3102.73 | 93159.0 | 47124.4 | - | - |
|  |  |  |  |  |  |  |  |  | [108.6; 131971.6] | [108.6; 55127.0] | - | - |
| East | AMG | 9 | -918.72 | 1855.45 | 8.86 | 0.01 | 775.93 | 5915.95 | 12292.9 | 4330.7 | 25.8 | 96.3 |
|  |  |  |  |  |  |  |  |  | [207.1; 827979.7] | [207.1; 232806.3] | [7.0; 44.6] | [53.6; 139.0] |
| Webster | AM2m | 10 | -510.09 | 1040.18 | - | 0.74 | 125.21 | 3837.23 | 432166.7 | 1567970.7 | - | - |
|  |  |  |  |  |  |  |  |  | [432166.7; 432166.7] | [1567633.8; 1568307.6] | - | - |
| Webster | IMG | 8 | -513.78 | 1043.55 | 3.38 | 0.14 | 121.13 | 3712.25 | 1386634.4 | 264957.2 | 0.2 | 9.9 |
|  |  |  |  |  |  |  |  |  | [1372001.6; 1401267.2] | [148792.5; 381122.0] | [0.0; 1.4] | [0.0; 24.6] |
| Webster | IM | 6 | -517.00 | 1046.01 | 5.83 | 0.04 | 119.97 | 3676.57 | 455767.2 | 1432599.8 | - | - |
|  |  |  |  |  |  |  |  |  | [452848.5; 458685.9] | [1425538.1; 1439661.6] | - | - |
| Webster | IM2NG | 10 | -513.24 | 1046.47 | 6.30 | 0.03 | 127.36 | 3903.28 | 695909.3 | 174486.3 | 0.5 | 27.1 |
|  |  |  |  |  |  |  |  |  | [648867.1; 742951.5] | [115410.6; 233562.0] | [0.0; 4.5] | [18.4; 35.7] |
| Webster | SIG | 6 | -518.14 | 1048.27 | 8.09 | 0.01 | 119.45 | 3660.85 | 387673.3 | 378214.1 | 2.0 | 39.9 |
|  |  |  |  |  |  |  |  |  | [386316.3; 389030.4] | [362835.0; 393593.2] | [1.6; 2.3] | [37.3; 42.5] |
| Webster | SI2NG | 8 | -516.20 | 1048.41 | 8.23 | 0.01 | 129.62 | 3972.43 | 746073.5 | 267102.4 | 0.9 | 64.7 |
|  |  |  |  |  |  |  |  |  | [740020.1; 752126.8] | [260992.5; 273212.2] | [0.5; 1.2] | [64.0; 65.4] |
| Webster | IM2mG | 11 | -513.76 | 1049.52 | 9.35 | 0.01 | 127.30 | 3901.23 | 1444451.6 | 229073.5 | 0.2 | 29.4 |
|  |  |  |  |  |  |  |  |  | [1422965.2; 1465938.0] | [148184.6; 309962.4] | [0.0; 2.1] | [17.8; 41.0] |
| Webster | SC2NG | 11 | -513.99 | 1049.98 | 9.81 | 0.01 | 114.12 | 3497.43 | 911342.4 | 308342.2 | 0.4 | 11.7 |
|  |  |  |  |  |  |  |  |  | [899950.7; 922734.1] | [300249.2; 316435.2] | [0.0; 1.7] | [10.8; 12.7] |
| Indian | SC2N2mG | 14 | -1089.03 | 2206.07 | - | 0.94 | 158.61 | 933.35 | 2048.1 | 957.3 | 64.0 | 6.3 |
|  |  |  |  |  |  |  |  |  | [1984.1; 2112.1] | [32.7; 8971.6] | [62.1; 65.9] | [4.1; 8.6] |
| Indian | SC2NG | 11 | -1094.92 | 2211.85 | 5.78 | 0.05 | 1201.86 | 7072.67 | 5306.7 | 4851.4 | 43.2 | 4.4 |
|  |  |  |  |  |  |  |  |  | [247.5; 709290.9] | [247.5; 1753775.2] | [42.4; 44.0] | [0.0; 77.7] |
| Cliff | SC2N2mG | 14 | -1006.59 | 2039.18 | - | 0.53 | 123.06 | 1482.41 | 21376.3 | 4560.6 | 24.7 | 19.6 |
|  |  |  |  |  |  |  |  |  | [9640.8; 33111.8] | [1982.4; 7138.8] | [20.9; 28.5] | [16.9; 22.3] |
| Cliff | SC2mG | 12 | -1007.72 | 2039.43 | 0.25 | 0.46 | 232.67 | 2802.70 | 71589.0 | 9480.8 | 3.6 | 10.7 |
|  |  |  |  |  |  |  |  |  | [98.1; 519641.2] | [98.1; 118734.2] | [0.0; 132.8] | [0.0; 86.9] |
| LAKE | MODEL | k | hrf | $m_{e12}$ | $m_{e21}$ | $m_{e*12}$ | $m_{e*21}$ | $Tsplit$ | $Tpost-split$ | $P$ | $Q$ | $O$ |
| Témiscouata | SC2mG | 12 | - | 0.000 | 0.000 | 0.000 | 0.000 | 29996.49 | 7246.04 | 0.93 | - | 0.99 |
|  |  |  | - | [0.000; 0.005] | [0.000; 0.005] | [0.000; 0.001] | [0.000; 0.001] | [0.00; 180578.93] | [0.00; 48478.87] | [0.22; 0.95] | - | [0.99; 0.99] |
| Témiscouata | SIG | 6 | - | - | - | - | - | 29506.96 | - | - | - | 0.99 |
|  |  |  | - | - | - | - | - | [48741.01; 31157.00] | - | - | - | [0.84; 0.99] |
| East | SC2N2mG | 14 | 0.11 | 0.020 | 0.040 | 0.002 | 0.001 | 35907.02 | 19601.58 | 0.60 | 0.16 | 0.98 |
|  |  |  | [0.01; 0.56] | [0.008; 0.032] | [0.017; 0.064] | [0.000; 0.008] | [0.000; 0.006] | [0.00; 85224.95] | [10542.92; 28660.24] | [0.26; 0.94] | [0.01; 0.37] | [0.96; 0.98] |
| East | IM2mG | 11 | - | 0.003 | 0.000 | 0.000 | 0.001 | 14597.50 | - | 0.77 | - | 0.97 |
|  |  |  | - | [0.000; 0.004] | [0.000; 0.001] | [0.000; 0.000] | [0.001; 0.001] | [0.00; 33933.42] | - | [0.59; 0.95] | - | [0.94; 0.99] |
| East | AM2N2m | 12 | 0.04 | 0.015 | 0.001 | 0.000 | 0.003 | 4204.02 | 0.00 | 0.06 | 0.50 | 0.96 |
|  |  |  | [0.01; 0.34] | [0.008; 0.021] | [0.000; 0.007] | [0.000; 0.002] | [0.000; 0.007] | [3263.59; 7077.67] | [0.00; 2730.24] | [0.00; 0.86] | [0.26; 0.74] | [0.96; 0.96] |
| East | AMG | 9 | - | 0.003 | 0.000 | - | - | 8725.63 | 1536.75 | - | - | 0.96 |
|  |  |  | - | [0.000; 0.007] | [0.000; 0.000] | - | - | [4276.72; 14127.18] | [0.00; 6368.43] | - | - | [0.95; 0.97] |
| Webster | AM2m | 10 | - | 0.001 | 0.008 | 0.000 | 0.000 | 35653.78 | 0.00 | 0.05 | - | 0.99 |
|  |  |  | - | [0.001; 0.001] | [0.000; 0.008] | [0.000; 0.001] | [0.000; 0.000] | [35401.86; 35905.70] | [0.00; 65494.31] | [0.01; 0.09] | - | [0.98; 0.99] |
| Webster | IMG | 8 | - | 0.000 | 0.000 | - | - | 35415.16 | - | - | - | 0.99 |
|  |  |  | - | [0.000; 0.001] | [0.000; 0.000] | - | - | [31876.76; 38953.56] | - | - | - | [0.99; 0.99] |
| Webster | IM | 6 | - | 0.000 | 0.000 | - | - | 35990.39 | - | - | - | 0.99 |
|  |  |  | - | [0.000; 0.001] | [0.000; 0.000] | - | - | [32383.78; 39596.99] | - | - | - | [0.98; 0.99] |
| Webster | IM2NG | 10 | 0.28 | 0.000 | 0.000 | - | - | 34499.01 | - | - | 0.01 | 0.99 |
|  |  |  | [0.08; 0.47] | [0.000; 0.001] | [0.000; 0.001] | - | - | [29327.19; 39670.83] | - | - | [0.00; 0.03] | [0.98; 0.99] |
| Webster | SIG | 6 | - | - | - | - | - | 37893.82 | - | - | - | 0.99 |
|  |  |  | - | - | - | - | - | [33197.67; 42589.97] | - | - | - | [0.98; 0.99] |
| Webster | SI2NG | 8 | 0.53 | - | - | - | - | 36252.68 | - | - | 0.04 | 0.99 |
|  |  |  | [0.09; 0.96] | - | - | - | - | [30137.19; 42368.17] | - | - | [0.02; 0.06] | [0.99; 0.99] |
| Webster | IM2mG | 11 | - | 0.000 | 0.000 | 0.000 | 0.000 | 35073.11 | - | 0.21 | - | 0.99 |
|  |  |  | - | [0.000; 0.001] | [0.000; 0.000] | [0.000; 0.000] | [0.000; 0.000] | [32731.26; 37414.95] | - | [0.01; 0.95] | - | [0.99; 0.99] |
| Webster | SC2NG | 11 | 0.59 | 0.000 | 0.000 | - | - | 4805.63 | 31892.69 | - | 0.04 | 0.99 |
|  |  |  | [0.47; 0.72] | [0.000; 0.001] | [0.000; 0.001] | - | - | [0.00; 23478.82] | [28130.88; 35654.50] | - | [0.01; 0.50] | [0.98; 0.99] |
| Indian | SC2N2mG | 14 | 0.20 | 0.002 | 0.004 | 0.000 | 0.002 | 29401.99 | 8541.27 | 0.60 | 0.41 | 0.97 |
|  |  |  | [0.01; 0.43] | [0.000; 0.003] | [0.002; 0.005] | [0.000; 0.002] | [0.002; 0.002] | [23833.27; 34970.72] | [6279.02; 10803.53] | [0.19; 0.95] | [0.39; 0.42] | [0.94; 1.00] |
| Indian | SC2NG | 11 | 0.19 | 0.000 | 0.001 | - | - | 6230.14 | 16705.74 | - | 0.50 | 0.95 |
|  |  |  | [0.01; 0.38] | [0.000; 0.011] | [0.000; 0.012] | - | - | [5259.77; 7200.51] | [0.00; 396069.76] | - | [0.48; 0.50] | [0.29; 0.99] |
| Cliff | SC2N2mG | 14 | 0.17 | 0.000 | 0.000 | 0.000 | 0.000 | 32375.74 | 9245.76 | 0.48 | 0.40 | 0.99 |
|  |  |  | [0.00; 1.00] | [0.000; 0.001] | [0.000; 0.002] | [0.000; 0.001] | [0.000; 0.009] | [28981.58; 35769.90] | [8722.07; 9769.46] | [0.36; 0.61] | [0.00; 1.00] | [0.98; 0.99] |
| Cliff | SC2mG | 12 | - | 0.009 | 0.022 | 0.000 | 0.000 | 12818.87 | 9679.91 | 0.05 | - | 0.96 |
|  |  |  | - | [0.000; 0.022] | [0.009; 0.034] | [0.000; 0.002] | [0.000; 0.001] | [2190.82; 23446.92] | [0.00; 19505.76] | [0.00; 0.26] | - | [0.95; 0.97] |

### Inference of model parameters

The inferred proportion of correctly oriented markers in the unfolded JAFS (parameter O) ranged from 95.4% to 99%, suggesting that the vast majority of ancestral allelic states were correctly inferred using the European Whitefish as an outgroup (Table 1, Supplementary Table S1). Considering only the highest ranked model for each lake, some general patterns emerged from the comparisons of inferred model parameters among lakes. First, differences in effective population sizes between dwarf and normal whitefish were inferred after splitting from the ancestral population in all lakes except East Lake (ANOVA, Témiscouata, *p* = 0.016; East, *p* = 0.67; Webster, *p* = 2.10^-12^; Indian, *p* = 1.13x10^-4^ and Cliff, *p* = 1.78x10^-7^, Table 1, Supplementary Table S1). When such differences were observed, *N_e_* was generally larger for dwarf compared to normal whitefish, except in Webster Lake. Taking into account population growth in four of the five lakes (N_e_ was inferred constant in Webster) revealed quite similar patterns of temporal population size changes, with recent demographic expansions being found in all populations except for the normal population of Témiscouata. A more pronounced demographic expansion was generally inferred for dwarf compared to normal whitefish (in Témiscouata, East and Indian lakes), and contemporary effective population size was also larger in dwarf than in normal populations in the four lakes where a secondary contact was inferred (ANOVA, Témiscouata, *p* = 0.095., marginally significant p<0.1; East, *p* = 0.047*; Indian, *p* = 0.04* and Cliff, *p* = 0.02*).

Asymmetric migration rates were also found in all five lakes for the two categories of loci assumed in heterogeneous migration models (−2m) (Table 1, Supplementary Table S1). The best models for the three most divergent species pairs (Webster, Indian and Cliff lakes) involved similar asymmetric migration directions for both categories of loci, with higher rates from dwarf to normal whitefish (m_e21_ > m_e12_ and m_e_'_21_ > m_e_'_12_) in Webster and Indian lakes and the opposite in Cliff Lake. In these three lakes, the proportion of loci exhibiting reduced effective migration rates generally followed the divergence continuum (Webster: 0.05; Indian: 0.41; Cliff: 0.52).The contemporary number of migrants exchanged per generation from one population to the other, estimated using the weighted mean effective migration rate in each direction (*i.e.,* average gene flow = *N*b*(P*m*_e_+(1-*P*)* *m*_e'_)), revealed a more pronounced gene flow from dwarf to normal populations in all lakes, except in Cliff Lake (ANOVA, Témiscouata, *p* = 9x10^-4^; East, *p* = 4x10^-4^; Webster, *p* = 1.99x10^-9^ and Indian, *p* = 0.023) (Fig. 5).

**Figure 5:**
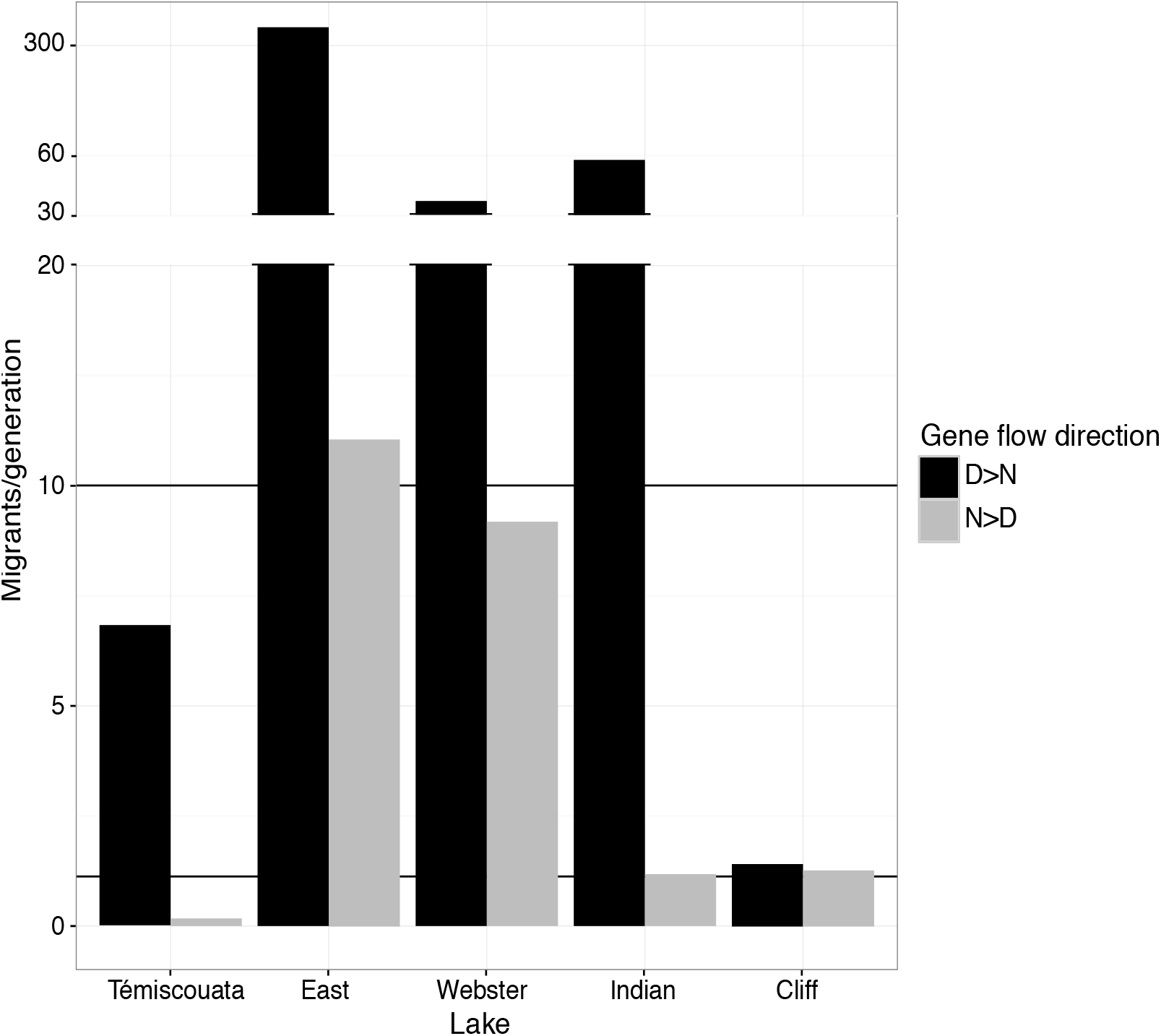
Asymmetrical effective gene flow between species among lakes. Bar plot of the number of migrants per generation for both directions from dwarf to normal (black) and reciprocally (gray), obtained from estimated parameters of the fittest model for each lake, using the average-gene flow formula *(i.e.,* average-gene flow = N*b*(P*m_e_+(1-P)* m_e_) in each direction.

The highest ranked model for East Lake and the two most divergent pairs (Indian and Cliff) included heterogeneous effective population size at the genomic level (Fig. 1C, Table 1). The fraction of the genome with a reduced *N*e (Q) was estimated to about 16% in East Lake, and 40% in Indian and Cliff lakes, and the degree of reduction in *N_e_ (i.e.,* the Hill-Robertson factor, *hrf)* was about 11%, 20% and 6% for East, Indian and Cliff lakes, respectively.

Time parameters, namely the duration of allopatric isolation (T_S_) and gene flow *(T_SC_* and *T_AM_),* were converted into absolute time estimates (years). Summing over periods of strict isolation and gene flow revealed a recent and similar divergence history in all five lakes *(Ts* was approximately 30,000; 36,000; 36,000; 29,000; 32,000 yrs for Témiscouata, East, Webster, Indian and Cliff lakes respectively) (Table 1). The inferred time of secondary contact in Témiscouata, East, Indian and Cliff lakes coincided roughly with the last glacial retreat following the Wisconsinian glaciation between 18,000 and 11,000 yrs before present (7,200; 19,600; 8,500 and 9,200 ybp, respectively, Table 1).

### Comparisons of genetic variation among lakes

Of the 42,582 SNPs that were genotyped in total (all lakes combined), only 2.5% corresponded to shared polymorphic loci across all lakes *(i.e.,* 'shared across lakes' category; Fig. 1). Reciprocally, about 25% of the SNPs were private to Témiscouata or East lakes, whereas Webster, Indian and Cliff lakes each contained ∼10% of private SNPs. The remaining majority of the loci were segregating in at least two lakes (see percentages on Fig. 1). Over the five lakes, a higher proportion of the loci were private to normal populations, compared to the dwarf populations. Within lakes, the highest proportions of SNPs private to normal whitefish were found in Témiscouata (51%) and Webster (69%). The three other lakes displayed the opposite pattern with highest proportion of private loci to the dwarf populations. Shared variation within lakes represented only 6% to 16% of the SNPs.

Partitioning genetic variation within and among lakes using a dAPC revealed distinct signals along the four first axes (Fig. 6A). On the first axis (LD1, explaining 39.5% of the variance), populations clustered by lakes according to their geographical distribution, roughly separating the three southernmost lakes (Webster, Indian and Cliff, negative coordinates) from the other two lakes (Témiscouata and East, positive coordinates). The second axis (LD2, not shown here but see Fig. S2, explaining 22.5% of the variance) mostly separated dwarf and normal whitefish from Cliff. The third axis (LD3, explaining 16% of the variance) tended to separate species pairs by shifting normal whitefish of Mississippian/Atlantic origin towards positive coordinates and dwarf whitefish of Acadian origin towards negative coordinates, except for East Lake. The two most extreme populations values on that axis corresponded to normal species from Cliff Lake, the least introgressed relict of the Mississippian/Atlantic lineage (Gagnaire et al. 2013b), and the dwarf species from Cliff Lake associated with the Acadian lineage. Finally, the fourth axis (LD4, explaining 11% of the variance) separated the dwarf and normal species from each lake, thus illustrating the shared ancestry between species among lakes (Fig. 6B).

**Figure 6:**
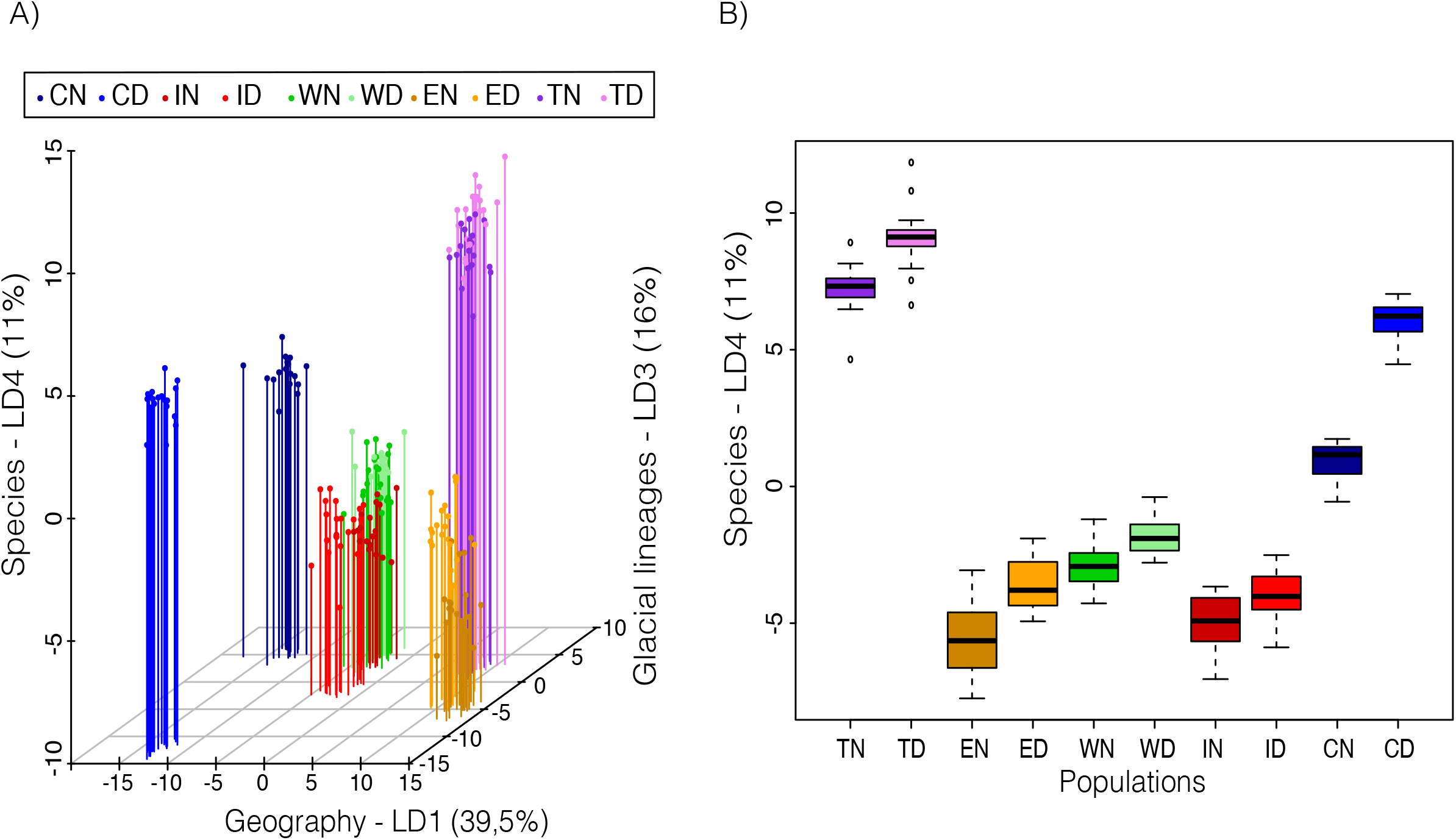
Genetic structure and relationship among lakes and species. **(A)** Discriminant analysis of principal components (dAPC) of the different lakes (Cliff, Indian, Webster, East, Témiscouata) for either Dwarf or Normal whitefish (D or N), presenting the three dimensional relationships between populations. First axis (LD1-39.5%), clustering of populations by lakes, corresponds to geographical distribution of the lakes. The LD3 (16%) separates species pairs and clusters dwarf and normal populations in two distinct groups associated to glacial lineages. Positive coordinates were occupied by normal populations of Atlantic origin and negative coordinates by dwarf populations of Acadian origin. The third axis, LD4 (11%) separated species within each lake. The LD2 is not shown here to avoid redundancy with LD1, which separated lakes populations. **(B)** Boxplot of the fourth dAPC axis (LD4), highlighting the divergence parallelism between species among lakes.

The genetic relationships among populations analyzed with *TreeMix* revealed two levels of signal (Fig. 7). The first level was directly linked with genetic distance between species. The population tree rooted with the normal population from Cliff (the most divergent population that best reflects the ancestral state of the Mississippian/Atlantic lineage, Lu et al. 2001) clearly separated normal whitefish from Cliff (CN) and Indian (IN) and dwarf whitefish from Cliff (CD) and Indian (ID), which were grouped together separately from all other populations. The clustering of CN with IN as that of CD and ID most likely reflect their shared ancestral polymorphism associated with their glacial lineage origin (Mississippian/Atlantic lineage for CN and IN; Acadian for CD and ID) (Lu et al. 2001). The second level of signal (geographic signal) grouped population pairs by lake in the remaining part of the tree, most likely reflecting the effect of gene flow following secondary contact between glacial lineages in each lake.

**Figure 7:**
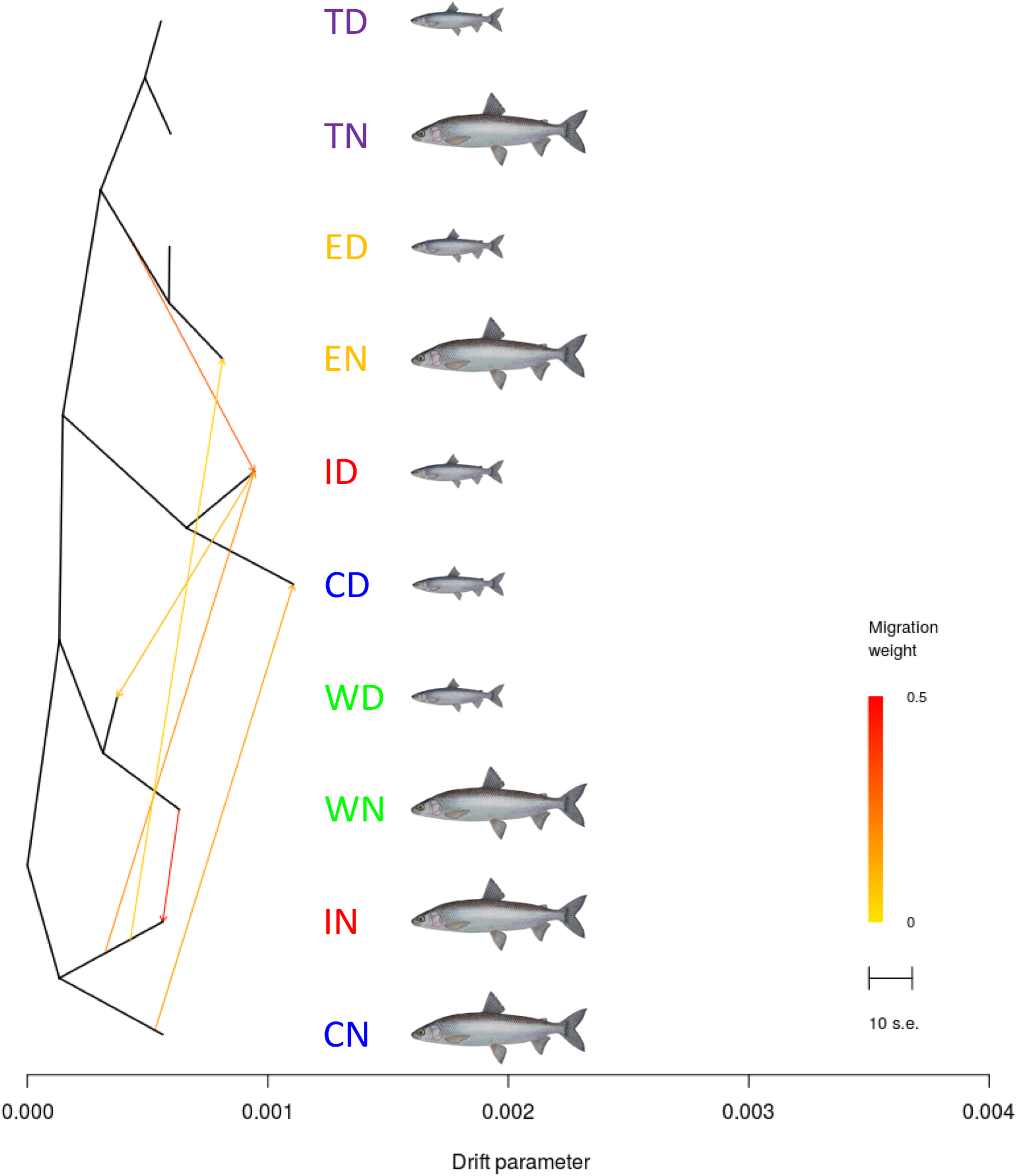
Shared ancestral genetic variation between allopatric populations and migration events between sympatric species pairs highlighted by genetic relationship among populations. The drift parameter axis is used as relative temporal measure where the scale bar indicates 10 times the average standard error (s.e.) of the relatedness among populations based on the variance-covariance matrix of allele frequencies. Color-scale indicates the inferred weight migration events.

Inferred migration links were represented by arrows, the color of which indicates their relative weights (Fig. 7). Migrations links between sympatric species pairs for Cliff and Indian lakes suggested contemporary gene flow *(i.e.,* consecutive to the colonization of the post-glacial lakes) between dwarf and normal populations within each of these two lakes. Other migration links between allopatric populations *(i.e.,* populations from different lakes which are isolated since the lakes formation) of the same species illustrated the genetic proximities *(i.e.* shared ancestry) of species from distinct lakes. For instance, the dwarf population from Webster (WD) was related to the dwarf population from Indian lake (ID), and the same link was found between the normal populations of these lakes (WN and IN). Finally, the ancestral population of East Lake was related with the dwarf population from Indian, whereas the normal population from East Lake (EN) was linked with the normal population from Indian Lake, thus supporting a common genetic background between the normal populations of East and Indian lakes.

### Discussion

Patterns of genomic differentiation among five dwarf/normal Lake Whitefish species pairs provide new insights into the divergence history of a well-studied model of ecological speciation (Bernatchez et al. 2010). The first approach implemented here relied on inferring the divergence history of each species pair separately, using the JAFS as a summary statistics of genome-wide differentiation patterns. In order to maximize the amount of available information, each JAFS was oriented using the closely related European whitefish as an outgroup species, thus providing increased power to detect demographic processes that generate asymmetric distributions of derived variants around the diagonal of the JAFS.

The secondary contact scenario was the most likely divergence history inferred for four of the five replicate species pairs. Moreover, this scenario was detected both in the least divergent (Témiscouata and East) and the most divergent (Indian and Cliff lakes) pairs. Therefore, our ability to detect the secondary contact was probably not affected by the small degree of genetic divergence between the least differentiated species pairs. Using simulations, Roux et al. (2016) recently showed that the power to detect the SC scenario can be high when the period of isolation is long relative to the duration of secondary contact. Moreover, secondary contacts are expected to leave detectable signatures on the JAFS, since the erosion of past allopatric divergence by secondary gene flow typically generates an excess of shared intermediate frequency alleles.

The secondary contact scenario is concordant with previous mtDNA-based phylogeograhic studies in northeastern America (Bernatchez and Dodson 1990; Bernatchez and Dodson 1991; Pigeon et al. 1997), and provided new insights into the evolutionary history of whitefish radiation. The geographic area where sympatric whitefish species pairs occur corresponds to a well-known suture zone where glacial lineages have come into contact in several freshwater species, as they were recolonizing from different refugia after the Laurentide ice sheet retreat (Curry 2007; April et al. 2013). In Lake Whitefish, this zone corresponds to a phylogeograhic transition between Acadian and Atlantic/Mississippian mitochondrial lineages (Bernatchez and Dodson 1990; Pigeon et al. 1997). Interestingly, the Allegash River basin (including the studied lakes), which represents the core of this contact zone, is the only area where sympatric populations of Lake Whitefish are observed. Moreover, no dwarf population, either in allopatry or sympatry, has been reported outside this region (Bernatchez and Dodson 1990). Therefore, phenotypic and ecological divergence, and in particular the occurrence of the dwarf phenotype, is tightly linked with the secondary contact zone.

The frequency of Acadian and Atlantic/Mississippian mitochondrial lineages within lakes was shown to be partly associated with level of phenotypic divergence between dwarf and normal whitefish, with variable amounts of mitochondrial introgression being found among lakes (Bernatchez and Dodson 1990; Pigeon et al. 1997). At one extreme, the least phenotypically divergent pair from East Lake is fixed for the Acadian mitochondrial haplogroup in both dwarf and normal whitefish, which has previously been taken as a support for sympatric divergence in this particular lake (Pigeon et al. 1997). Although our inferences based on the JAFS could not definitely rule out the IM model (IM2mG, *w_AIC_* = 0.19), we obtained much stronger evidence in favor of the secondary-contact scenario in this lake (SC2N2mG, *w_AIC_* = 0.77). A possible explanation for the loss of the Atlantic/Mississippian haplogroup in East Lake involves a stronger recent demographic expansion in the dwarf population following secondary contact, which likely contributed to the fixation of the Acadian lineage. Indeed, the preferential direction of introgression between hybridizing populations with asymmetrical *Ne* is expected to occur from the larger to the smaller population (Barton 1986). This is consistent with the preferential direction of the effective gene flow inferred here (Fig. 5). This interpretation is also supported by the closely similar scenario inferred in the neighboring Témiscouata Lake, which harbors the second least divergent species pair. Témiscouata Lake is also dominated by the Acadian haplogroup, but a small proportion of normal whitefish in this lake is still associated with the Atlantic/Mississippian lineage, indicating incomplete mitochondrial swamping. Since we also inferred an expansion of the dwarf population (but not in the normal whitefish) following secondary contact in this lake, it is likely that this demographic imbalance explains the predominance of Acadian mitochondrial haplotypes in the northern part of the contact zone. At the other extreme, Cliff Lake where species divergence is the most pronounced shows differential fixation of Acadian and Atlantic/Mississippian haplotypes in dwarf and normal populations, respectively (Bernatchez and Dodson 1990). Thus, there is a perfect association in this lake between glacial lineage origin and phenotypic divergence, which was also attributed to a secondary contact in our demographic inferences. Similarly to East Lake, Indian Lake harbored both dwarf and normal populations fixed for the Acadian haplogroup (Lu et al. 2001). However, our analysis of the JAFS also confirmed that two distinct glacial lineages have come into contact in this lake. Such a partial concordance is typically expected when secondary contact occurs between incompletely reproductively isolated species (Taylor and McPhail 2000).

Less clear results were obtained from our demographic inferences in Webster Lake. In this lake, both the normal and the dwarf populations display mitochondrial introgression (Pigeon et al. 1997). Consistent with bi-directional gene flow after secondary contact, the Acadian haplogroup is more frequent in the dwarf population, whereas the Atlantic/Mississippian is the most common lineage in the normal population (Pigeon et al. 1997). However, the JAFS was best explained by the AM2m model (w_AIC_ *=* 0.74). Webster Lake is known to have been particularly impacted by human activities since the early 1800's due to artificial connections established with a distinct watershed for water regulation. Therefore, we cannot exclude that contemporary whitefish populations from Webster Lake have been impacted by recent admixture events between distinct populations from different lakes. This recent history may have confounded the detection of the secondary contact in this lake, which nevertheless ranks among the best-retained models (SC2NG, ΔAIC < 10).

### A shared history of divergence before independent evolution within lakes

A global analysis including all five pairs simultaneously was necessary to understand the extent to which replicate whitefish species pairs share a common history of divergence. The secondary contact scenario implies that the different population pairs are derived from the same two glacial lineages, the evidence of which is partly supported by mitochondrial data (Bernatchez and Dodson 1990; Pigeon et al. 1997). However, whether whitefish species pairs share a common history before secondary contact has never been assessed using nuclear markers.

Grouping populations based on their overall genetic similarities with *Treemix* produced two different types of grouping in the population tree. Populations from the three least divergent species pairs were grouped by lake *(i.e.,* TN grouped with TD, EN with ED, and WN with WD), while populations from the two most divergent species pairs were grouped by ecotypes *(i.e.,* IN with CN, and ID with CD). This complex picture likely reflects the relative importance of gene flow between species within lakes and genetic drift among lakes, and is in itself insufficient to distinguish contemporary admixture from shared ancestry during lakes colonization. Inferring migration events among populations enabled us to detect current gene flow between sympatric divergent species pairs in Indian and Cliff lakes. However, the other inferred links connecting populations of the same species but from different lakes *(i.e.,* isolated populations) rather indicated shared genetic variation due to common ancestry. Namely, inferred links between Webster and Indian indicated the sharing of ancestral variation between WN and IN (and therefore with CN), as well as between WD and ID (and therefore with CD). This supports the view that the different populations of each species in these three lakes, which are not connected today by gene flow, were genetically similar before being isolated in their respective lakes. An additional link inferred between EN and IN (itself connected with WN and CN) confirmed that normal whitefish from East Lake share ancestral variation with other normal populations from the southern part of the contact zone. This provides further evidence that the secondary contact inferred in East Lake has occurred between the same two glacial lineages as for the other lakes, despite the lack of Atlantic/Mississippian mitochondrial lineage in this lake. Finally, the ancestral population from East Lake was linked to the dwarf population from Indian (and therefore to WD and CD), indicating that both populations from East Lake share much of the ancestral variation originating from the Acadian lineage. This is also consistent with the genetic swamping hypothesis proposed for explaining the lack of mitochondrial polymorphism in this lake. The analysis of overall diversity patterns performed with the dAPC (Fig. 6) was a complementary way to disentangle remaining signals of genetic differentiation between glacial lineages (axis 3) from genetic differentiation among lakes (axis 1). On the third axis, the projection of dwarf and normal populations from Cliff Lake indicated the positions of the two least introgressed populations of our dataset. Therefore, they could be used to define an Acadian (negative coordinates) and an Atlantic/Mississippian (positive coordinates) reference for comparisons with other lakes. Interestingly, both populations from East and Indian lakes occupied intermediate positions, which is concordant with a higher proportion of Acadian ancestry in these lakes, as suggested by mitochondrial data (Pigeon et al. 1997).

In summary, the most parsimonious overall scenario supported by our analyses corresponds to a secondary contact for all lakes, with variable contributions of Acadian and Atlantic/Mississippian lineages due to demographic contingencies. The secondary contact was concomitant to population expansions in both glacial lineages, which were detected for most lakes. This is broadly consistent with the idea that the two glacial lineages were undergoing spatial expansions after the last glacial retreat, which provoked a secondary contact at the origin of parallel genetic divergence patterns across whitefish species pairs. Our results also support that population expansions were generally more pronounced for dwarf relative to normal populations, still reflected today by the higher contemporary abundance of dwarf whitefish in all lakes (L. Bernatchez, unpubl. data). This demographic imbalance also impacted the main direction of gene flow, which was more pronounced from dwarf to normal populations than the reverse. As a consequence, an important amount of shared ancestral polymorphism between dwarf and normal populations (Fig. 2) should correspond to genetic variation coming from gene exchanges due to introgression between lineages, in addition to incomplete lineage sorting.

### An extended framework for inferring speciation-with-gene-flow

The concept of speciation-with-gene-flow embraces a large diversity of divergence scenarios with regards to the timing of gene flow, which in turn pertains to different modes of speciation that have long been recognized in the speciation literature (Coyne and Orr 2004). Diverging populations can experience temporal variations in effective size and migration rate, which both influence the temporal dynamics of gene flow. Consequently, demographic inference methods that account for these temporal variations have the potential to provide an improved underpinning of the historical demographic events that shaped the unfolding of speciation.

For the Lake Whitefish as for other species with a pan-Arctic distribution, the history of divergence has been strongly impacted by quaternary climatic oscillations (Bernatchez and Wilson 1998). Glaciations have drastically restricted species distribution areas provoking geographic isolation among bottlenecked populations (Bernatchez et al. 1989; Ambrose 1998; Aoki et al. 2008), while inter-glacial periods have allowed secondary contacts between populations expanding from their glacial refugia (Hewitt 2001). Here, accounting for temporal variation in migration rate and *N_e_* allowed us to determine that the secondary contact between whitefish glacial lineages has occurred contemporarily with population expansions. This later point is of prime importance for understanding the evolution of reproductive isolation, since bottlenecked populations undergoing demographic expansions are likely to fix deleterious alleles (Luikart et al. 1998; Peischl et al. 2013; Lohmueller 2014), which could later translate into substrate for genetic incompatibilities cumulated during allopatric phase, upon secondary contact. Moreover, these genetic incompatibilities may associate, by coupling, to form barriers to gene flow (Barton and de Cara 2009; Bierne et al. 2011), as proposed in Gagnaire et al. (2013b).

Another important aspect of divergence-with-gene-flow relates to the extent to which the previously described demographic effects interact with selection. The speciation genomics literature is increasingly integrating the influence of selective processes in historical divergence models (e.g., Roux et al. 2013; Sousa and Hey 2013; Tine et al. 2014; Roux et al. 2016), and more generally, in the analytical approaches to relate genomic divergence patterns to the underling evolutionary processes (Cruickshank and Hahn 2014). These selective effects can be separated in two broad categories. First, genetic barriers caused by local adaptation and reproductive isolation loci can resist introgression, hence reducing the effective migration rate at linked loci (Barton and Bengtsson 1986; Feder and Nosil 2010). The second category embraces the effect of positive (Smith and Haigh 1974) and background selection (Charlesworth et al. 1993), which cause local reductions in genetic diversity at both selected sites and linked neutral sites. These later selective effects rather correspond to a reduction in the *Ne* of the genomic regions influenced by selection, irrespective to the role that they play in the speciation process. Since gene flow depends both on *Ne* and migration rate, both types of selective effects are likely to impact genomic divergence patterns during speciation. Here, we captured these effects separately using divergence-with-gene-flow models that take into account in a simple way the effects of genetic barriers and linked selection.

Accounting for variation in effective migration rate across the genome generally improved the fits to empirical data whatever the model considered (Fig. 4B), and the best models for all lakes also included heterogeneous migration rates. This suggests that the rate of introgression between whitefish glacial lineages has been highly variable across their genome since the beginning of secondary contact, as reported previously in other species (Tine et al. 2014; Le Moan et al. 2016; Rougemont et al. 2016). Moreover, integrating heterogeneous *N_e_* in the models also improved model scores for the two most divergent species pairs (Cliff and Indian, Fig. 4C). Therefore, our results also support the view that linked selection has influenced the patterns of genomic divergence in whitefish. As proposed in earlier studies, this mechanism may be particularly efficient in low-recombining chromosomal regions (Cruickshank and Hahn 2014). Some of our models (SC2m2N and AM2m2N) combined both genome-wide variation in *Ne* and *me,* as already developed within an ABC framework (Roux et al. 2016). The rationale behind this is that only models that both contain a period of isolation and gene flow enable to dissociate the influence of both sources of chromosomal variation, since only linked selection is at play during periods of geographic isolation. However, it is currently unclear how much the signal contained in empirical polymorphism data can retain distinct signatures for these two selective effects. This will need to be addressed using simulations.

In sum, our approach illustrates the need to take into account both temporal and genomic variations in effective population size and migration rates when inferring the history of speciation. The 26 divergence models considered here enabled us to evaluate a large diversity of scenarios, taking each effect separately an in combination with other to improve the inference of the divergence history while controlling for model complexity.

### Understanding the divergence continuum in whitefish

Lake Whitefish nascent species pairs offer a rare opportunity to understand the influence of selection and historical demography on a continuum of phenotypic and genomic divergence associated with speciation. Previous works have provided mounting evidence for the role of selection in shaping genetic and phenotypic divergence across this continuum (Rogers et al. 2002; Landry et al. 2007; Bernatchez et al. 2010; Renaut et al. 2012; Gagnaire et al. 2013b; Laporte et al. 2015). However, the role played by historical demography has never been fully resolved since previous studies largely depended on mitochondrial DNA alone.

Our study brings new evidence supporting previous findings based on mitochondrial DNA that the onset of this young adaptive radiation matched the last glacial period (Bernatchez and Dodson 1990; Bernatchez and Dodson 1991). Using a mutation rate of 10^-8^ mutations/site/generation and a generation time of 3.5 years, the average divergence time between glacial lineages was 41,600 years (s.d. 8,100). This is consistent with Jacobsen et al. (2012), who estimated the divergence time between 20,000 and 60,000 ybp based on mitogenome sequencing. This corresponds to the late Wisconsin glacial episode; starting 85,000 years ago, during which the Laurentide ice sheet covered the studied region in eastern North America, with a maximum ice extent occurring around 25,000 ybp (Curry 2007). The average time of secondary contact obtained here, was dated to 11,200 years (s.d. 5,700), which also corresponds to the glacial retreat period, at which the lakes were colonized by the two glacial lineages from eastern (Acadian) and western (Atlantic/Mississipian) glacial refuges (again, 18,000 to 11,000 ybp) as described by Curry (2007). Therefore, the inferred timing of divergence and secondary contact between glacial lineages matches relatively well the chronology of the climatic events in eastern North America.

Our results suggest that demographic differences among lakes have contributed to shaping the divergence continuum observed among the five lakes. Introgression rates tended to be higher in the least divergent species pairs, resulting into a weaker genetic differentiation. Yeaman et al. (2016) recently showed that the formation of genomic islands by erosion of divergence following secondary contact depends on the amount of linkage disequilibrium (LD) among selected loci and the intensity of effective migration. Here, we showed that effective migration rate was generally higher in the least differentiated lakes (Témiscouata, East and Webster), while at the same time, increased LD among islands has been documented in the most divergent lakes (Gagnaire et al. 2013b). Therefore, the divergence continuum likely implies both the antagonistic effects of divergent selection maintaining LD and introgression eroding past divergence.

Our study also provides new insights on the role of linked selection in shaping patterns of genomic divergence observed among the whitefish species pairs. Namely, we inferred that some genomic regions have experienced a reduction in *N_e_,* as predicted under the effect of selection at linked sites (Cruickshank and Hahn 2014). The increasing proportion of genomic regions affected by Hill-Robertson effects, from the less to the most divergent lakes, indicated that the divergence continuum among lakes was also influenced by linked selection. In the light of those observations, along with previous studies, we propose that the continuum of genetic divergence in whitefish species pairs is the evolutionary result of a complex interplay between *(i)* genetic divergence between glacial lineages through lineage sorting and mutation accumulation, *(ii)* reduced introgression in genomic regions involved in reproductive isolation due to the accumulation of incompatibilities, (iii) divergent selection on phenotypes maintaining LD, and (iv) the independent contingency of demographic events among lakes. The heterogeneous landscape of species divergence in the whitefish system was thus likely built by a combination of selective and demographic factors. Our inferences allowed us to disentangle part of this complex interplay, although many aspects remain to be clarified. In particular, whether selection at linked sites also plays a role in facilitating the accumulation of incompatibilities during allopatry will need to be scrutinized into more details, as well as the role of such incompatibilities in facilitating the divergence of quantitative polygenic traits after contact. This could be done by testing the effect of divergence on quantitative traits with and without the joint action of selection against hybrids. Indeed, a model mixing components of allopatric speciation, with the accumulation of genetic incompatibilities (e.g., under-dominant mutations) and sympatric speciation *(i.e.,* local adaptation involving divergent selection on quantitative traits), would differ from the coupling hypothesis model which mostly considers local adaptation loci of relatively strong effect (Bierne et al. 2011). We argue that this kind of models could be relevant for some systems in which sympatric speciation after admixture, or parallel hybrid speciation, has been inferred without explicitly testing a single divergence event with recent secondary gene flow (Meier et al. 2016; Kautt et al. 2016). We also believe that demographic inferences approaches should systematically include basic scenario of divergence, extended by models with increasing levels of complexity to address demographic and selective effects separately and then combined, and not only focus on the *a priori* history of the system. To conclude, this study provides a clear illustration of the potential benefits to apply the models developed here towards disentangling the relative role of selective and demographic processes towards elucidating the complexity of species divergence in any other taxonomic group.

## Materials and Methods

### Sampling and genotyping

We used RAD-sequencing data from Gagnaire et al. (2013b) to generate a new genome-wide polymorphism dataset. Previous studies based on these data (Gagnaire et al. 2013b; Laporte et al. 2015) only focused on a subset of 3438 RAD markers that were included in the Lake Whitefish linkage map (Gagnaire et al. 2013a). Here, we used the total amount of sequence data (1.7* 10^9^ reads of 101 bp) to document genome-wide variation in five sympatric species pairs occurring in five isolated lakes from the Saint-John River basin (Fig. 1). For each pair, 20 normal and 20 dwarf individuals were used for RAD-sequencing, but five individuals that received poor sequencing coverage were removed from the dataset. Consequently, the following analyses were performed with 195 individuals, each having an average number of 8.7x10^6^ of reads.

We also sequenced six European Whitefish *(Coregonus lavaretus),* a sister species closely related to the North American Lake Whitefish, to provide an outgroup for identifying ancestral and derived alleles at each polymorphic site within the American Lake Whitefish. European Whitefish were sampled in Skrukkebukta Lake (Norway, 69°34'11.6”N-30°02'31.9”E), which also harbors postglacial sympatric whitefish species pairs (Amundsen et al. 2004; Østbye et al. 2006). RAD libraries were prepared for three individuals from each ecotype population, using the same procedure as for American Lake Whitefish (Gagnaire et al. 2013b).

The *C. lavaretus* raw sequence dataset was filtered using the same criteria as for *C. clupeaformis* (Gagnaire et al. 2013b). After sequence de-multiplexing, the reads were trimmed to a length of 80 bp to avoid sequencing errors due to decreasing data quality near the end of reads. We then used the *Stacks* pipeline (v1.24) for *de novo* RAD-tags assembly and individual genotyping (Catchen et al. 2013). We used a custom *Perl* script to determine an optimal set of assembly parameters for *Ustacks.* A minimal coverage depth of 5x per allele (m=5) and a maximal number of six mismatches between two haplotypes at a same locus within individuals (M=6) were set. We then allowed a maximal number of six mismatches between individuals in *Cstacks* (n=6) to merge homologous RAD-tags from different samples. Finally, we used the program *Population* to export a VCF file containing the genotypes of all individuals.

Several filtering steps were then performed with *VCFtools* v0.1.13 (Danecek et al. 2011) to remove miscalled and low-quality SNPs, as well as false variation induced by the merging of paralogous loci. We first removed SNPs with more than 10% missing genotypes in each on the 10 *C. clupeaformis* populations. A lower exclusion threshold of 50% was used for *C. lavaretus* to retain a maximum of orthologous loci with the outgroup. We then filtered for Hardy-Weinberg disequilibrium within each of the 10 *C. clupeaformis* populations, using a p-value exclusion threshold of 0.01. Finally, we merged the filtered datasets of dwarf and normal populations within each lake together with the European whitefish outgroup and kept only loci that passed the previous filters in all three samples. This resulted in five lake-outgroup datasets containing 14812, 22788, 5482, 26149, and 14452 SNPs for Témiscouata, East, Webster, Indian and Cliff lakes, respectively. Variation in SNP number for Webster Lake was associated to a lower coverage that reduced the amount of detectable markers and also an overall reduced level of polymorphism in that lake. Finally, we determined the most parsimonious ancestral allelic state for loci that were monomorphic in the outgroup but polymorphic in *C. clupeaformis.* The resulting oriented SNP datasets contained 11985, 11315, 5080, 13905 and 9686 SNPs for Témiscouata, East, Webster, Indian and Cliff lakes respectively, that were used to build the unfolded JAFS of each lake, using customs *perl* and *R* scripts. In each lake, the JAFS was projected down to 13 individuals *(i.e.,* 26 chromosomes) per population to avoid remaining missing genotypes and optimize the resolution of the JAFS.

### Inferring the history of divergence with gene flow

Because the five lakes are isolated from each other since around 12,000 years (Curry 2007), we analyzed their JAFS separately. The demographic and selective histories of the five species pairs were inferred using a custom version of the software *dadi* v1.7 (Gutenkunst et al. 2009). We considered 26 models (Fig. 2) that were built to extend four basic models representing alternative modes of divergence: Strict Isolation (SI), Isolation-with-Migration (IM), Ancient Migration (AM), and Secondary Contact (SC). Each model consists of an ancestral population of size N_ref_ that splits into two populations of effective size N_1_ and N_2_ during T_S_ (SI, IM), T_AM_+T_S_ (AM), or T_S_+T_SC_ (SC) generations, possibly exchanging migrants during T_S_ (IM), T_AM_ (AM), or T_SC_ (SC) generations at rate m_e12_ from population 2 *(i.e.,* normal populations) into population 1 *(i.e.,* dwarf populations), and m_e21_ in the opposite direction. These models were extended to integrate temporal variation in effective population size (-G) by enabling exponential growth using current-to-ancient population size ratio parameters *b*1 (for dwarf populations) and *b*2 (for normal populations) to account for expansions or bottlenecks. Variation in effective population size across the genome due to Hill-Robertson effects (Hill and Robertson 1966) - *i.e.* local reduction in *N_e_* at linked neutral sites due to the effect of background (Charlesworth et al. 1993) and positive selection (Smith and Haigh 1974) - was modeled by considering two categories of loci (-2N) occurring in proportions *Q* and 1*-Q.* In order to quantify a mean effect of selection at linked sites, we defined a Hill-Robertson scaling factor *(hrf),* relating the effective population size of loci influenced by selection *(N'_1_=hrf×N_1_* and N'_2_=hrf×N_2_) to that of neutral loci (N_1_ and N_2_). Then, models of divergence with gene flow were extended to account for heterogeneous migration across the genome by considering two categories of loci (-2m). In addition to a first category of loci evolving neutrally *(i.e.,* with migration rates m_e12_ and m_e21_) and occurring in proportion *P,* we considered a second category of loci that occur in proportion 1-P, experiencing different effective migration rates m_e_'_12_ and m_e_'_21_ due to their linkage with nearby selected genes (Tine et al. 2014). Because migration and drift influence gene flow during the whole divergence time in the IM model, the effects of heterogeneous migration and population effective size were evaluated separately (IM2N vs. IM2m). However, these effects could be estimated jointly in AM and SC models using the period without gene flow to decouple the effects of migration and drift (-2N2m in addition to −2N and −2m). All models with heterogeneous gene flow were also implemented to allow for population growth (−2NG, −2mG and −2N2mG). Finally, in order to take into account potential errors in the identification of ancestral allelic states, predicted JAFS were constructed using a mixture of correctly oriented and mis-orientated SNPs occurring in proportions *O* and 1-O, respectively.

The 26 models were fitted independently for each lake using successively a hot and a cold simulated annealing procedure followed by 'BFGS' optimization (Tine et al. 2014). We ran 25 independent optimizations for each model in order to check for convergence and retained the best one (Supplementary Table S2) to perform comparisons among models based on Akaike information criterion (AIC). Our comparative framework thus addresses overparametrization issues by penalizing models which contain more parameters. By allowing comparisons among nested models of increasing complexity, it also provides a means to independently evaluate the effect of accounting for temporal or genomic variation in migration rate or effective population size. A conservative threshold was applied to retain models with *ΔAIC_i_ = AIC_i_ — AIC_min_ <* 10, since the level of empirical support for a given model with a Δ*AIC_i_* > 10 is essentially none (Burnham and Anderson 2002). For each lake, the difference in AIC between the worst and the best model *Δ_max_ = AIC_max_ — AIC_min_* was used to obtain a scaled score for each model using:

(1)

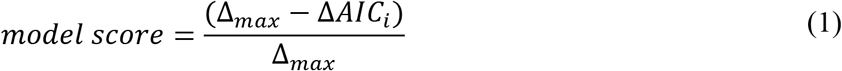

such that for each lake the worst model takes a score of 0 and the best model takes a score of 1. Therefore, the model score could be used to more easily compare the retained models among lakes. Akaike weights (w_AIC_) were also computed following equation (2) to evaluate the different models within each lake, where *R* corresponds to the number of models *(R* = 26).

(2)

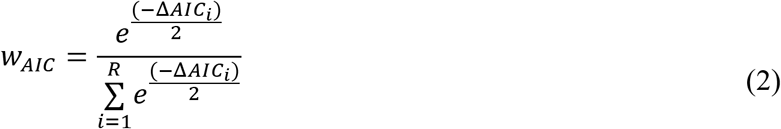

Finally, we converted estimated demographic parameters into biologically meaningful units in order to compare informative parameter values among lakes (e.g. timing and strength of gene flow). These estimates were only used for indication and comparative purpose, since we were missing crucial information about the per generation mutation rate in Lake Whitefish. We used the optimal multiplicative scaling factor *theta* between model and data to estimate the ancestral effective population size *(N*ref) before split for each lake:

(3)

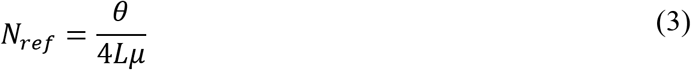

with *μ* being the mutation rate (fixed at 10” mutations/site/generation) and *L* the effective length of the genome explored by our RAD-Seq experiment and estimated as:

(4)

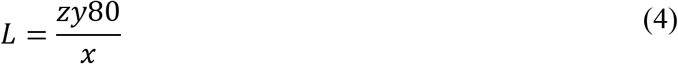

where x is the number of SNPs that were originally detected from *y* RAD-tags of 80 bp present in the initial dataset, and *z* is the number of SNP retained for *∂a∂i* analyses in the lake considered. Estimated times in units of 2*N*ref generations were converted into years assuming a generation time of 3.5 years *(i.e.,* the average between 3 years for dwarf and 4 years for normal whitefish), and estimated migration rates were divided by 2*N*ref to obtain the proportion of migrants received by each population every generation.

### Patterns of shared ancestry and admixture among lakes

Because the analyses of historical divergence were performed separately in each species pair, we also searched for signatures of shared ancestry and gene flow among replicate species pairs to provide a broader understanding of the divergence history in whitefish. The five lake-specific datasets used for demographic inferences were merged together and only polymorphic loci that were retained after filtering in all five lakes were considered (42558 SNPs in total). For each lake, we determined the fraction of private polymorphisms, the fraction of SNPs shared with at least one other lake or shared across all five lakes. We then measured the proportion of SNPs that were shared between species and private to each species within each lake, as well as among lakes.

To visualize the overall genetic structure and relationships among lakes and species, we performed a discriminant analysis of principal components (dAPC) in *Adegenet* v2.0.0 (Jombart et al. 2010). We first imputed missing genotypes within each population using a Random Forest regression approach, which provides a more accurate imputation than the replacement of missing genotypes by population mean allele frequency (Poland et al. 2012). Imputation was performed using ten iterations with 150 trees using the *randomForestSRC* v1.6.1 package in *Stackr* v.0.1.5 (Gosselin and Bernatchez 2016), and imputed sub-datasets were subsequently merged to perform the dAPC.

Finally, we used *TreeMix* v1.12 (Pickrell and Pritchard 2012) to infer historical relationships among populations. This method uses the covariance structure of allele frequencies between populations and a Gaussian approximation for genetic drift to build a maximum likelihood graph relating sampled populations to their common ancestor, taking migration events into account to improve the fit to the inferred tree. Migration events between populations are modeled in *TreeMix* by discrete mixture events. Such events may either reflect gene exchange between populations within lakes and/or genetic correlations between geographically isolated populations of the same species, due to the retention of shared ancestral polymorphism among populations from different lakes following their geographic isolation. In order to avoid interpreting spurious migration signals, we focused on the main events of gene flow that received the highest weights, which likely correspond to the largest admixture proportions. We thus allowed a maximum of six migration events to be inferred among branches of the whitefish population tree. For this analysis, we used a 20% missing genotype rate per population without imputing missing genotypes to avoid potential biases in the covariance matrix.

## Acknowledgements

We would like to thank Nicolas Bierne, Anne-Laure Ferchaud, Martin Laporte, Anne-Marie Dion-Côté and Charles Perrier for insightful discussions, Thierry Gosselin for *stackr* inputs, as well as Anne C. Dalziel for commenting an earlier version of this manuscript. We are grateful to Guillaume Côté, Melissa L. Evans, William Adam and the staff of Maine Department of Inland Fisheries & Wildlife (David J. Basley and Jeremiah Wood) for all of their help sampling whitefish and sharing information about watersheds histories, and to Kim Præbel for *Coregonus lavaretus* RAD-seq raw data. This research was supported by a discovery research grant from the Natural Sciences and Engineering Research Council of Canada (NSERC) to L.B. L.B also holds the Canadian Research Chair in genomics and conservation of aquatic resources, which funded the research infrastructure for this project.

## References

Ambrose SH. 1998. Late Pleistocene human population bottlenecks, volcanic winter, and differentiation of modern humans. Journal of Human Evolution 34:623-651.

Amundsen P-A, Bøhn T, Vâga GH. 2004. Gill raker morphology and feeding ecology of two sympatric morphs of European whitefish (Coregonus lavaretus). Annales Zoologici Fennici:291-300.

Aoki K, Kato M, Murakami N. 2008. Glacial bottleneck and postglacial recolonization of a seed parasitic weevil, Curculio hilgendorfi, inferred from mitochondrial DNA variation. Molecular Ecology 17:3276-3289.

April J, Hanner RH, Dion-Côté A-M, Bernatchez L. 2013. Glacial cycles as an allopatric speciation pump in north-eastern American freshwater fishes. Molecular Ecology 22:409-422.

Avise JC. 2000. Phylogeography: the history and formation of species.

Barton NH, Bengtsson BO. 1986. The barrier to genetic exchange between hybridising populations. Heredity 57:357-376.

Barton NH. 1986. The effects of linkage and density-dependent regulation on gene flow. Heredity 57: 415-426.

Barton NH, de Cara MAR. 2009. The evolution of strong reproductive isolation. Evolution 63:1171-1190.

Becquet C, Przeworski M. 2007. A new approach to estimate parameters of speciation models with application to apes. Genome Research 17:1505-1519.

Bernatchez L, Dodson JJ, Boivin S. 1989. Population bottlenecks: influence on mitochondrial DNA diversity and its effect in coregonine stock discrimination. Journal of Fish Biology 35:233-244.

Bernatchez L, Dodson JJ. 1990. Allopatric origin of sympatric populations of lake whitefish (Coregonus clupeaformis) as revealed by mitochondrial-DNA restriction analysis. Evolution 24:890-908.

Bernatchez L, Dodson JJ. 1991. Phylogeographic structure in mitochondrial DNA of the lake whitefish (Coregonus clupeaformis) and its relation to Pleistocene glaciations. Evolution 45:1016-1035.

Bernatchez L, Renaut S, Whiteley AR, Derome N, Jeukens J, Landry L, Lu G, Nolte AW, Ostbye K, Rogers SM, et al. 2010. On the origin of species: insights from the ecological genomics of lake whitefish. Philos. Trans. R. Soc. Lond., B, Biol. Sci. 365:1783-1800.

Bernatchez L, Wilson CC. 1998. Comparative phylogeography of Nearctic and Palearctic fishes. Molecular Ecology 7:431-452.

Bernatchez L. 2004. Ecological theory of adaptive radiation: an empirical assessment from coregonine fishes (Salmoniformes). Pages 175-207 in A.P. Hendry and S.C. Stearns, eds. Evolution illuminated: salmon and their relatives. Oxford Univ. Press, Oxford.

Bierne N, Welch J, Loire E, Bonhomme F, David P. 2011. The coupling hypothesis: why genome scans may fail to map local adaptation genes. Molecular Ecology 20:2044-2072.

Burnham KP, Anderson DR. 2002. Model selection and multimodel inference: a practical information-theoretic approach.

Butlin RK, Saura M, Charrier G, Jackson B, André C, Caballero A, Coyne JA, Galindo J, Grahame JW, Hollander J, et al. 2014. Parallel evolution of local adaptation and reproductive isolation in the face of gene flow. Evolution 68:935-949.

Catchen J, Hohenlohe PA, Bassham S, Amores A, Cresko WA. 2013. Stacks: an analysis tool set for population genomics. Molecular Ecology 22:3124-3140.

Charlesworth B, Morgan MT, Charlesworth D. 1993. The effect of deleterious mutations on neutral molecular variation. Genetics 134:1289-1303.

Charlesworth B, Nordborg M, Charlesworth D. 1997. The effects of local selection, balanced polymorphism and background selection on equilibrium patterns of genetic diversity in subdivided populations. Genet. Res. 70:155-174.

Charlesworth D. 2006. Balancing Selection and Its Effects on Sequences in Nearby Genome Regions. PLOS Genetics 2(4): e64. doi:0.1371/journal.pgen.0020064

Corbett-Detig RB, Hartl DL, Sackton TB. 2015. Natural Selection Constrains Neutral Diversity across A Wide Range of Species.Barton NH, editor. PLoS Biol 13:e1002112.

Cornuet J-M, Pudlo P, Veyssier J, Dehne-Garcia A, Gautier M, Leblois R, Marin J-M, Estoup A. 2014. DIYABC v2.0: a software to make approximate Bayesian computation inferences about population history using single nucleotide polymorphism, DNA sequence and microsatellite data. Bioinformatics 30:1187-1189.

Coyne JA, Orr HA. 2004. Speciation. Sunderland MA Sinauer Associates Inc

Cruickshank TE, Hahn MW. 2014. Reanalysis suggests that genomic islands of speciation are due to reduced diversity, not reduced gene flow. Molecular Ecology 23:3133-3157.

Curry RA. 2007. Late glacial impacts on dispersal and colonization of Atlantic Canada and Maine by freshwater fishes. Quaternary Research 67:225-233.

Danecek P, Auton A, Abecasis G, Albers CA, Banks E, DePristo MA, Handsaker RE, Lunter G, Marth GT, Sherry ST, et al. 2011. The variant call format and VCFtools. Bioinformatics 27:2156-2158.

Duvaux L, Belkhir K, Boulesteix M, Boursot P. 2011. Isolation and gene flow: inferring the speciation history of European house mice. Molecular Ecology 20:5248-5264.

Ewing GB, Jensen JD. 2016. The consequences of not accounting for background selection in demographic inference. Molecular Ecology 25:135-141.

Excoffier L, Dupanloup I, Huerta-SÃ nchez E, Sousa VC, Foll M. 2013. Robust Demographic Inference from Genomic and SNP Data. Akey JM, editor. PLoS Genet 9:e1003905-e1003917.

Feder JL, Egan SP, Nosil P. 2012. The genomics of speciation-with-gene-flow. Trends in Genetics 28:342-350.

Feder JL, Nosil P. 2010. The efficacy of divergence hitchhiking in generating genomic islands during ecological speciation. Evolution 64:1729-1747.

Gagnaire P-A, Normandeau E, Pavey SA, Bernatchez L. 2013a. Mapping phenotypic, expression and transmission ratio distortion QTL using RAD markers in the Lake Whitefish (Coregonus clupeaformis). Molecular Ecology 22:3036-3048.

Gagnaire P-A, Pavey SA, Normandeau E, Bernatchez L. 2013b. The genetic architecture of reproductive isolation during speciation-with-gene-flow in lake whitefish species pairs assessed by RAD sequencing. Evolution 67:2483-2497.

Gosselin T, Bernatchez L. 2016. stackr: GBS/RAD Data Exploration, Manipulation and Visualization using R. R package version 0.2.1. https://github.com/thierrygosselin/stackr.

Gutenkunst RN, Hernandez RD, Williamson SH, Bustamante CD. 2009. Inferring the joint demographic history of multiple populations from multidimensional SNP frequency data. McVean G, editor. PLoS Genet 5:e1000695

Harrison RG. 1990. Hybrid zones: windows on evolutionary process. Oxford surveys in evolutionary biology 7:69-128.

Harrison RG, Larson EL. 2016. Heterogeneous genome divergence, differential introgression, and the origin and structure of hybrid zones. Molecular Ecology 25:2454-2466.

Hewitt G. 2000. The genetic legacy of the Quaternary ice ages. Nature 405:907-913.

Hewitt GM. 1996. Some genetic consequences of ice ages, and their role in divergence and speciation. Biological Journal of the Linnean Society 58:247-276.

Hewitt GM. 2001. Speciation, hybrid zones and phylogeography—or seeing genes in space and time. Molecular Ecology 10:537-549.

Hewitt GM. 2004. Genetic consequences of climatic oscillations in the Quaternary. Philosophical Transactions of the Royal Society of London B: Biological Sciences 359:183-95-discussion195.

Hey J, Nielsen R. 2004. Multilocus methods for estimating population sizes, migration rates and divergence time, with applications to the divergence of Drosophila pseudoobscura and D. *persimilis*. Genetics 167:747-760.

Hey J, Nielsen R. 2007. Integration within the Felsenstein Equation for Improved Markov Chain Monte Carlo Methods in Population Genetics. Proc. Natl. Acad. Sci. U.S.A. 104:2785-2790.

Hill WG, Robertson A. 1966. The effect of linkage on limits to artificial selection. Genet. Res. 8:269-294.

Jacobsen, M. W., M. M. Hansen, L. Orlando, D. Bekkevold, L. Bernatchez, E. Willerslev, and M. T. P. Gilbert. 2012. Mitogenome sequencing reveals shallow evolutionary histories and recent divergence time between morphologically and ecologically distinct European whitefish (Coregonus spp.). Molecular Ecology 21:2727-2742.

Jombart T, Devillard S, Balloux F. 2010. Discriminant analysis of principal components: a new method for the analysis of genetically structured populations. BMC Genet. 11:94.

Kautt AF, Machado-Schiaffino G, Meyer A. 2016. Multispecies Outcomes of Sympatric Speciation after Admixture with the Source Population in Two Radiations of Nicaraguan Crater Lake Cichlids.Payseur BA, editor. PLoS Genet 12:e1006157-33.

Landry L, Bernatchez L. 2010. Role of epibenthic resource opportunities in the parallel evolution of lake whitefish species pairs (Coregonus sp.). J. Evol. Biol. 23:2602-2613.

Landry L, Vincent WF, Bernatchez L. 2007. Parallel evolution of lake whitefish dwarf ecotypes in association with limnological features of their adaptive landscape. J. Evol. Biol. 20:971-984.

Laporte M, Rogers SM, Dion-Côté A-M, Normandeau E, Gagnaire P-A, Dalziel AC, Chebib J, Bernatchez L. 2015. RAD-QTL Mapping Reveals Both Genome-Level Parallelism and Different Genetic Architecture Underlying the Evolution of Body Shape in Lake Whitefish (Coregonus clupeaformis) Species Pairs. G3 (Bethesda) 5:1481-1491.

Le Moan A, Gagnaire PA, Bonhomme F. 2016. Parallel genetic divergence among coastal-marine ecotype pairs of European anchovy explained by differential introgression after secondary contact. Molecular Ecology 25:3187-3202.

Lohmueller KE. 2014. The impact of population demography and selection on the genetic architecture of complex traits. Williams SM, editor. PLoS Genet 10:e1004379.

Lu G, Basley DJ, Bernatchez L. 2001. Contrasting patterns of mitochondrial DNA and microsatellite introgressive hybridization between lineages of lake whitefish (Coregonus clupeaformis); relevance for …. Molecular Ecology 10:965-985.

Lu G, Bernatchez L. 1999. Correlated trophic specialization and genetic divergence in sympatric lake whitefish ecotypes (Coregonus clupeaformis): support for the ecological speciation hypothesis. Evolution 53:1491-1505.

Luikart G, Allendorf FW, Cornuet JM, Sherwin WB. 1998. Distortion of allele frequency distributions provides a test for recent population bottlenecks. Journal of Heredity 89:238-247.

McPhail JD. 1992. Ecology and evolution of sympatric sticklebacks (Gasterosteus): evidence for a species-pair in Paxton Lake, Texada Island, British Columbia. Canadian Journal of Zoology 70:361-369.

Meier JI, Sousa VC, Marques DA, Selz OM, Wagner CE, Excoffier L, Seehausen O. 2016. Demographic modeling with whole genome data reveals parallel origin of similar Pundamilia cichlid species after hybridization. Molecular Ecology.

Nachman MW, Payseur BA. 2011. Recombination rate variation and speciation: theoretical predictions and empirical results from rabbits and mice. Philosophical Transactions of the Royal Society B: Biological Sciences 367:409-421.

Nosil, P. 2012. Ecological speciation. Oxford Univ. Press, Oxford, U.K

Nosil P. 2008. Speciation with gene flow could be common. Molecular Ecology 17:2103-2106.

Payseur BA. 2010. Using differential introgression in hybrid zones to identify genomic regions involved in speciation. Mol Ecol Resour 10:806-820.

Peischl S, Dupanloup I, Kirkpatrick M, Excoffier L. 2013. On the accumulation of deleterious mutations during range expansions. Molecular Ecology 22:5972-5982.

Pickrell JK, Pritchard JK. 2012. Inference of population splits and mixtures from genome-wide allele frequency data. PLoS Genet 8:e1002967.

Pigeon D, Chouinard A, Bernatchez L. 1997. Multiple Modes of Speciation Involved in the Parallel Evolution of Sympatric Morphotypes of Lake Whitefish (Coregonus clupeaformis, Salmonidae). Evolution 51:196.

Pinho C, Hey J. 2010. Divergence with gene flow: models and data. Annual review of ecology 41:215-230.

Poland J, Endelman J, Dawson J, Rutkoski J, Wu S, Manes Y, Dreisigacker S, Crossa J, Sánchez-Villeda H, Sorrells M, et al. 2012. Genomic Selection in Wheat Breeding using Genotyping-by-Sequencing. The Plant Genome Journal 5:103-111.

Renaut S, Maillet N, Normandeau E, Sauvage C, Derome N, Rogers SM, Bernatchez L. 2012. Genome-wide patterns of divergence during speciation: the lake whitefish case study. Philos. Trans. R. Soc. Lond., B, Biol. Sci. 367:354-363.

Rogers SM, Bernatchez L. 2007. The genetic architecture of ecological speciation and the association with signatures of selection in natural lake whitefish (Coregonus sp. Salmonidae) species pairs. Molecular Biology and Evolution 24:1423-1438.

Rogers SM, Gagnon V, Bernatchez L. 2002. Genetically based phenotype-environment association for swimming behavior in lake whitefish ecotypes (Coregonus clupeaformis Mitchill). Evolution 56:2322-2329.

Rougemont Q, Gagnaire P-A, Perrier C, Genthon C, Besnard A-L, Launey S, Evanno G. 2016. Inferring the demographic history underlying parallel genomic divergence among pairs of parasitic and nonparasitic lamprey ecotypes. Molecular Ecology doi: 10.1111/mec.13664.

Roux C, Fraïsse C, Romiguier J, Anciaux Y, Galtier N, Bierne N. 2016. Shedding light on the grey zone of speciation along a continuum of genomic divergence. bioRxiv doi: http://dx.doi.org/10.1101/059790

Roux C, Tsagkogeorga G, Bierne N, Galtier N. 2013. Crossing the Species Barrier: Genomic Hotspots of Introgression between Two Highly Divergent Ciona intestinalis Species. Molecular Biology and Evolution 30:1574-1587.

Schluter D. 1996. Ecological Speciation in Postglacial Fishes. Philosophical Transactions of the Royal Society of London B: Biological Sciences 351:807-814.

Smith JM, Haigh J. 1974. The hitch-hiking effect of a favourable gene. Genet. Res. 23:23-35.

Sousa V, Hey J. 2013. Understanding the origin of species with genome-scale data: modelling gene flow. Nature Publishing Group 14:404-414.

Swenson NG, Howard DJ. 2005. Clustering of Contact Zones, Hybrid Zones, and Phylogeographic Breaks in North America. Am Nat 166:581-591.

Taylor EB, Bentzen P. 1993. Evidence for Multiple Origins and Sympatric Divergence of Trophic Ecotypes of Smelt (Osmerus) in Northeastern North America. Evolution 47:813.

Taylor EB. 1999. Species pairs of north temperate freshwater fishes: evolution, taxonomy, and conservation. Reviews in Fish Biology and Fisheries 9:299-324.

Taylor EB, Donald McPhail J. 2000. Historical contingency and ecological determinism interact to prime speciation in sticklebacks, Gasterosteus. Proceedings of the Royal Society B: Biological Sciences 267:2375-2384.

Tine M, Kuhl H, Gagnaire P-A, Louro B, Desmarais E, Martins RST, Hecht J, Knaust F, Belkhir K, Klages S, et al. 2014. European sea bass genome and its variation provide insights into adaptation to euryhalinity and speciation. Nature Communications 5:1-10.

Welch JJ, Jiggins CD. 2014. Standing and flowing: the complex origins of adaptive variation. Molecular Ecology 23:3935-3937.

Wolf JBW, Ellegren H. 2016. Making sense of genomic islands of differentiation in light of speciation. Nature Reviews Genetics:1-14.

Wood CC, Foote CJ. 1996. Evidence for Sympatric Genetic Divergence of Anadromous and Nonanadromous Morphs of Sockeye Salmon (Oncorhynchus nerka). Evolution 50:1265.

Wu C-I. 2001. The genic view of the process of speciation. J. Evol. Biol. 14:851-865.

Yeaman S, Aeschbacher S, Bürger R. 2016. The evolution of genomic islands by increased establishment probability of linked alleles.Abbott RJ, Barton NH, Good JM, editors. Molecular Ecology 25:2542-2558.

Østbye K, Amundsen P-A, Bernatchez L, Klemetsen A, Knudsen R, Kristoffersen R, Naesje TF, Hindar K. 2006. Parallel evolution of ecomorphological traits in the European whitefish Coregonus lavaretus (L.) species complex during postglacial times. Molecular Ecology 15:3983-4001.

